# Pseudouridylation landscape across 42 nuclear-encoded *S. cerevisiae* tRNAs

**DOI:** 10.64898/2026.04.28.721490

**Authors:** Margaret L. Barry, Robin L. Abu-Shumays, Lauren E. Barnes, Ethan A. Shaw, Julia L. Reinsch, Zachary D. Basham, Miten Jain, Kristin S. Koutmou, David M Garcia

**Affiliations:** Institute of Molecular Biology, University of Oregon, Eugene, OR 97403, USA; Department of Biology, University of Oregon, Eugene, OR 97403, USA; Biomolecular Engineering Department, University of California Santa Cruz, Santa Cruz, CA 95064, USA; Center for Molecular Biology of RNA, University of California Santa Cruz, Santa Cruz, CA 95064, USA; Department of Chemistry, University of Michigan, Ann Arbor, MI 48109, USA; Department of Bioengineering, Northeastern University, Boston, MA 02115, USA; Department of Physics, Northeastern University, Boston, MA 02115, USA; Khoury College of Computer Sciences, Northeastern University, Boston, MA 02115, USA

## Abstract

Pseudouridine is the most abundant RNA base modification due to its prevalence in tRNA and rRNA, where it serves as a key modulator of structure and function. Yet, in the widely used model organism, the budding yeast *Saccharomyces cerevisiae,* tRNA pseudouridine sites have not been comprehensively annotated. Here, we performed two high-throughput methods established for detecting tRNA pseudouridylation positions: Nanopore direct RNA sequencing (DRS) and 2-bromoacrylamide-assisted cyclization sequencing (BACS). Using DRS, we sequenced cytosolic tRNA from seven pseudouridine synthase (PUS) knockout *S. cerevisiae* strains, including deletion strains of Pus1, Pus3, and Pus7. Analysis of these data verified thirty-two existing pseudouridine annotations and predicted an additional ten previously unannotated pseudouridine sites. Our analysis also revealed DRS signal changes at several non-uridine sites with the loss of a PUS, including apparent changes in modification abundances at position 37 upon deletion of Pus3. Liquid chromatography tandem mass spectrometry (LC-MS/MS) and primer extension assays, however, indicated no change in the abundance of these modifications with the loss of Pus3, underscoring the need for caution in interpreting DRS-based signal changes. Using BACS, we confirmed the ten DRS-predicted novel sites, while further detecting eleven additional previously unannotated pseudouridine sites. Combining existing modification annotations from the Modomics database with our DRS and BACS datasets, we created a map of all detected pseudouridines—totaling 126 sites, including 21 novel sites—and the enzymes responsible for their catalysis, across the forty-two nuclear-encoded *S. cerevisiae* tRNA isoacceptors.

## INTRODUCTION

Transfer RNA (tRNA) is the most densely modified type of RNA, and pseudouridine (Ψ) is the most prevalent RNA modification (Fig. 1) (Rintala-Dempsey and Kothe 2017). Pseudouridine contributes to tRNA structure and stability, as well as translation and decoding (Schultz et al. 2024; Biela et al. 2025; Wu et al. 2025); reviewed in (Jackman and Alfonzo 2013; Grosjean and Westhof 2016). Despite its prevalence, pseudouridine has not been comprehensively mapped in the most well-studied eukaryotic model for tRNA modifications, the budding yeast *Saccharomyces cerevisiae*. In fact, 11/42 budding yeast cytosolic isoacceptors lack annotation of *any* modification in the Modomics database.

**Figure 1.**
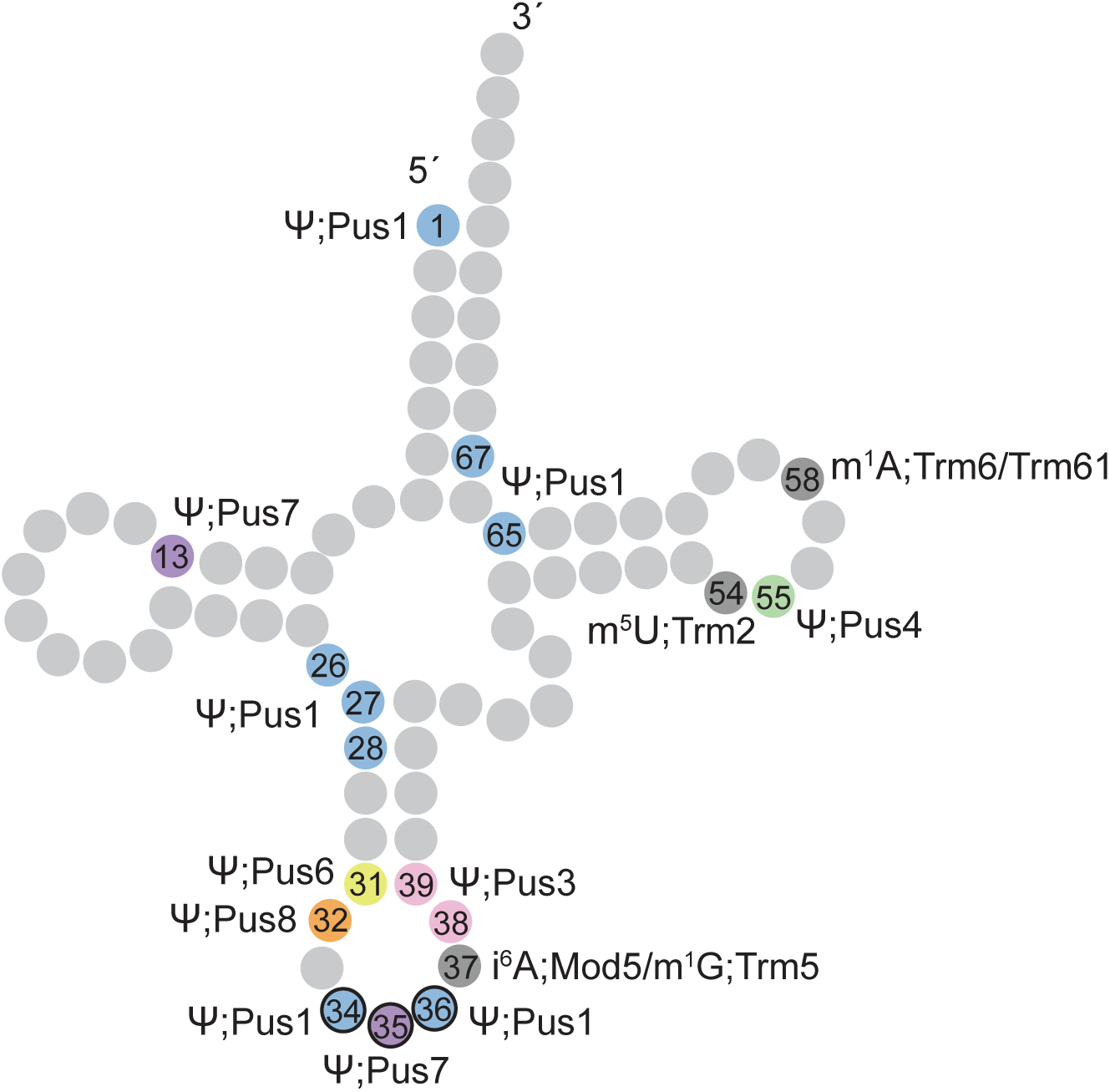
Generic *S. cerevisiae* nuclear-encoded tRNA annotated with known positions of pseudouridine (Ψ) and additional modifications discussed in this work. The catalyzing enzyme for each modification is denoted. Numbers indicate base position in the tRNA structural alignment. Base colors are specific to the catalyzing PUS enzyme. Pseudouridine sites match those shown in Figure 6. Non-Ψ modifications relevant to this study are colored dark gray; N6-isopentenyladenosine (i^6^A), 1-methylguanosine (m^1^G), 5-methyluridine (m^5^U), and 1-methyladenosine (m^1^A). Illustration is adapted from (Rintala-Dempsey and Kothe 2017).

*S.cerevisiae* is an established model for investigating tRNA modifications, as many modifications found in budding yeast are conserved across eukaryotes (reviewed in Phizicky and Hopper 2023). In some cases, addition of one modification impacts the addition of other modifications. Such interdependencies are known as modification “circuits” (reviewed in (Porat 2025). Emblematic of this, addition of pseudouridine to position 55 (Ψ_55_) has been shown to promote addition of both 5-methyluridine to position 54 (m^5^U_54_) and 1-methyladenosine to position 58 (m^1^A_58_) (Barraud et al. 2019; Yared et al. 2023; Shaw et al. 2024; Reinsch and Garcia 2026).

Oxford Nanopore Technologies direct RNA sequencing (DRS) has emerged as a valuable tool for profiling tRNA modifications, with pseudouridine among the most clearly detectable (Thomas et al. 2021; Lucas et al. 2024; Sun et al. 2023; White et al. 2023; Shaw et al. 2024; White et al. 2024, 2025; Rübsam et al. 2025; Riquelme-Barrios et al. 2025; Reinsch and Garcia 2026). Nanopore DRS presents the opportunity to profile the modification landscape of multiple tRNA isoacceptors across their entire sequence simultaneously. However, it currently lacks the ability to directly identify and quantify modifications. Most applications of Nanopore DRS for tRNA have relied on analysis of base miscalls, though recent work has moved toward direct analysis of ionic current (Rübsam et al. 2025; Akeson et al. 2025). Even modifications that are robustly detected using basecalling methods can present challenges in specific sequence contexts (Fleming et al. 2021, 2023; Liu-Wei et al. 2024; Rübsam et al. 2025). Liquid chromatography tandem mass spectrometry (LC-MS/MS) is valuable for its ability to unambiguously report the chemical identity of a modified nucleoside. When applied to samples digested down to individual nucleosides, it can be used to quantify the abundance of many chemical modifications simultaneously (Jones et al. 2023a). Position- and modification-specific assays that rely on reverse transcription (RT) stops or mutational signatures, caused either by modifications themselves or by modification-specific derivatives, can yield site-specific information including relative changes in modification abundance (Jackman et al. 2003; Zhang et al. 2019; Behrens et al. 2021; Zhang et al. 2022; Draycott et al. 2023; Xu et al. 2024; Hwang et al. 2025; Reinsch and Garcia 2026). These RT-based assays, however, can be complicated by non-specific reactions involving other modifications, particularly in densely modified molecules like tRNA. Thus, combining multiple methods permits a more comprehensive analysis of the positions and identities of tRNA chemical modifications.

We previously profiled the modification landscape of budding yeast nuclear-encoded tRNAs using Nanopore DRS coupled with LC-MS/MS (Shaw et al. 2024). Sequencing a genetic knockout of the <u>p</u>seudo<u>u</u>ridine <u>s</u>ynthase (PUS) Pus4 demonstrated pseudouridylation of U_55_ in nearly all cytosolic tRNAs. To complete the characterization of the pseudouridine landscape in these tRNAs, we applied DRS to additional mutant strains lacking an individual PUS: *pus1*Δ, *pus3*Δ, *pus6*Δ, and *pus7*Δ. To complement this approach, we performed total nucleoside LC-MS/MS, and the pseudouridine-specific detection method 2-bromoacrylamide-assisted cyclization sequencing (BACS) on wild-type, *pus1*Δ, *pus3*Δ, and *pus7*Δ strains. Our results identified both known and previously unannotated pseudouridine modifications. In combination with our previous data from a *pus4*Δ strain (Shaw et al. 2024), we created an annotated map of all known PUS-catalyzed *S. cerevisiae* nuclear-encoded tRNA pseudouridines. This includes 21 pseudouridines currently unannotated in the RNA modification database Modomics (Cappannini et al. 2024). We additionally provided examples of robust changes in DRS signal at positions coincident with other modifications that failed to be corroborated by orthogonal methods, emphasizing the need for caution in distinguishing true changes from sequencing artifacts.

## RESULTS

### Nanopore DRS and BACS identify both known and unknown Pus1 target sites

Pus1 localizes to the nucleus, recognizing structural motifs to modify tRNA, as well as mRNA and U2 snRNA (Simos et al. 1996; Motorin et al. 1998; Massenet et al. 1999; Arluison et al. 1999; Behm-Ansmant et al. 2006; Carlile et al. 2014; Grünberg et al. 2023). In budding yeast, Pus1 has been previously shown to modify multiple tRNAs at positions 1, 26, 27, 28, 34, 36, 65, and 67 among the 31 tRNA isoacceptors that have been annotated (Fig. 1) (Motorin et al. 1998; Arluison et al. 1999; Großhans et al. 2001; Behm-Ansmant et al. 2006; Khonsari and Klassen 2020; Cappannini et al. 2024). However, 11/42 budding yeast cytosolic isoacceptors currently lack annotation of any modifications, including pseudouridine, in the Modomics database.

To profile the modifications catalyzed by Pus1 across the 42 budding yeast cytosolic isoacceptors, we sequenced tRNA from wild-type and Pus1 deletion (*pus1*Δ) strains using Nanopore direct RNA sequencing (DRS). We then constructed heatmaps showing the probability that the base identity assigned from the Nanopore ionic current data matched the canonical reference base for that position (reference match probability) (Supplemental Fig. S1-2). This probability was calculated using an error model incorporating standard DRS mismatch rates, insertions, and deletions. The error model was derived from DRS of *in vitro* transcribed cytosolic tRNA sequences, and accounts for miscalls caused solely by sequence context (Shaw et al. 2024). Comparison of the probability of a miscall between wild-type and the knockout strain allowed visualization of the positions at which the ionic current signal had changed (Fig. 2A). Positions with a positive change in reference match probability indicate a decrease in miscalls in the knockout, potentially representing the loss of a modification. A negative change in value indicates an increase in miscalls, suggesting an increase in a modification in the knockout strain. We used an empirically determined change in reference match probability threshold value to identify all positions that changed with the loss of a given PUS enzyme. This threshold was calculated from the changes in reference match probability for PUS deletion strains (*pus1*Δ, *pus3*Δ, or *pus7*Δ) relative to wild-type, specifically at sites previously annotated to be modified by the corresponding enzyme (e.g., at U_27_ on tRNA^Arg(UCU)^ for Pus1) (See Materials and Methods and Supplemental Tables S1-S2). This threshold was then applied across all positions on all tRNAs to search for sites *de novo*.

**Figure 2.**
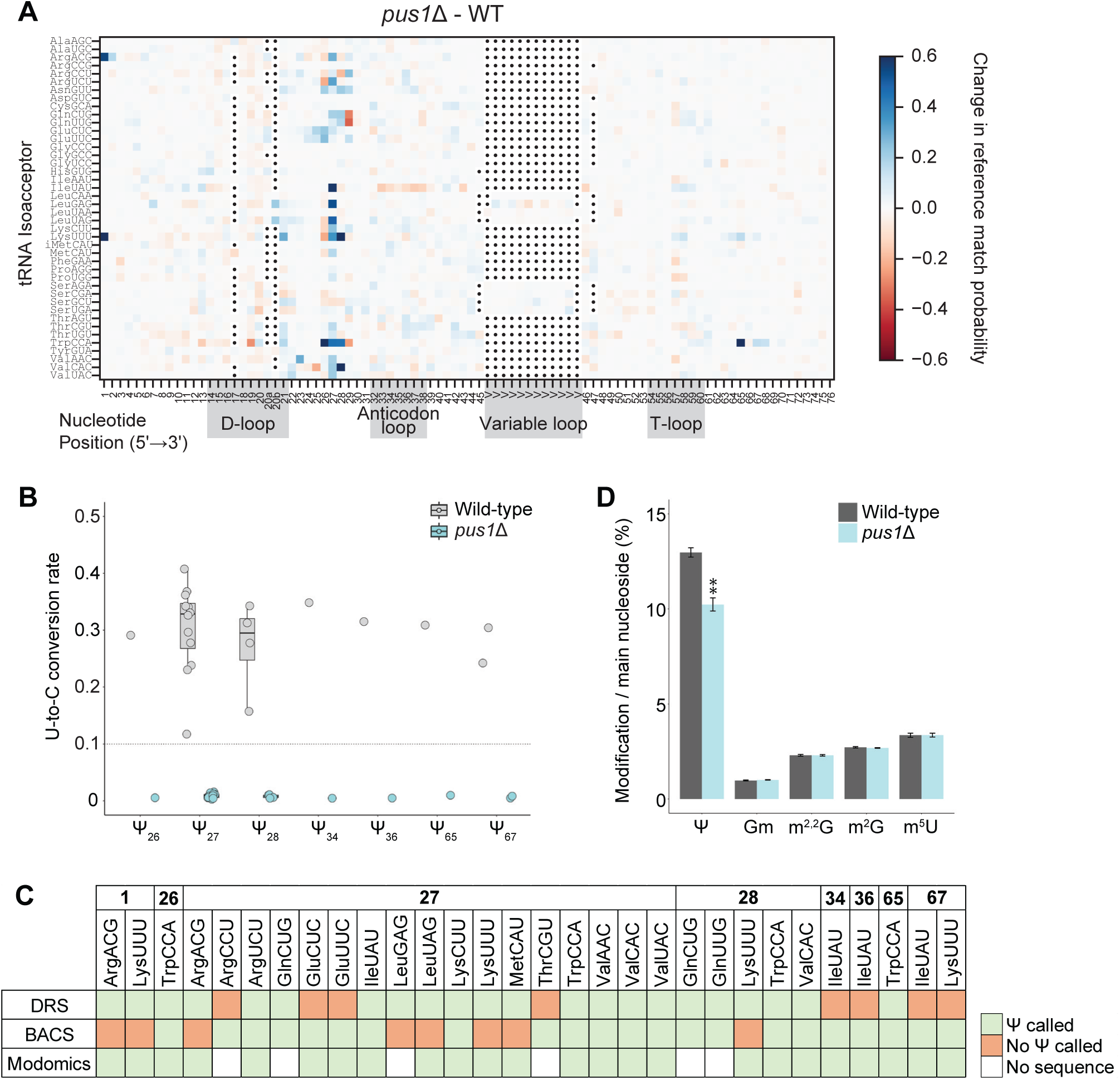
Modification of *pus1*Δ and wild-type *S. cerevisiae* nuclear-encoded tRNA. (A) Subtractive heatmap comparing Nanopore direct RNA sequencing (DRS) reference match probabilities of *pus1*Δ relative to wild-type. Reference match probability measures the probability that the DRS-called base matches the canonical base in the reference sequence. (B) U-to-C conversion rates from 2-bromoacrylamide-assisted cyclization sequencing (BACS) of wild-type and *pus1*Δ cytosolic tRNA. Each data point represents a single site on a single isoacceptor (e.g. Ψ_27_ on tRNA^Trp(CCA)^), averaged across three biological replicates. Boxplots show the 25% to 75% percentiles with a line at the median. Whiskers show 1.5 times the interquartile range. The conversion rate threshold for calling a pseudouridine (0.1) is marked with a dashed line. Only sites passing the threshold in the treated wild-type data are shown. *P* < 0.0001 for all sites, comparing wild-type and *pus1*Δ conversion rates (Šídák’s multiple comparisons test). Annotated or DRS-predicted pseudouridine sites that failed to pass the threshold with BACS are shown in Supplemental Figure S3A. (C) Summary of DRS and BACS-detected, and Modomics-annotated pseudouridine sites catalyzed by Pus1, grouped by tRNA position. “No sequence” refers to tRNAs for which no modifications are annotated in Modomics. (D) Total nucleoside LC-MS/MS of tRNA harvested from wild-type and *pus1*Δ strains. Modification abundances of pseudouridine (Ψ), 2’-O-methylguanosine (Gm), N2,N2-dimethylguanosine (m^2,2^G), N2-methylguanosine (m^2^G), and 5-methyluridine (m^5^U) are shown. Data for all measured modifications are available in Supplemental Table S6. “**” indicates *P* < 0.001 by Dunnett’s multiple comparison test of the mutant values compared to wild-type.

Comparing nanopore DRS data from *pus1*Δ and wild-type tRNA, we observed a decrease in miscalls in *pus1*Δ at eighteen sites that corresponded with Modomics-annotated Pus1 targets, i.e. specific uridines modified by Pus1 at specific positions in specific tRNA isoacceptors (Fig. 2A, Supplemental Tables S2&S3). Four additional sites occurring at positions known to be modified by Pus1 in other isoacceptors showed a signal change passing the threshold, potentially representing novel pseudouridine annotations (Supplemental Tables S2&S3).

To orthogonally validate DRS-predicted pseudouridines, we performed 2-bromoacrylamide-assisted cyclization sequencing (BACS) (Xu et al. 2024, 2025). Treatment of RNA with 2-bromoacrylamide causes pseudouridine to undergo chemical conversion leading to a Ψ-to-C mutational signature in reverse transcription. We performed BACS on tRNA from wild-type and *pus1*Δ cells and compared U-to-C conversion rates between the two samples (Fig. 2B, Supplemental Table S4). Three of the four previously unannotated Pus1 sites predicted by DRS showed high U-to-C conversion rates in the wild-type BACS data that were reduced to below-threshold levels in *pus1*Δ, indicating that these are true pseudouridine sites. BACS also identified two additional Pus1-catalyzed, unannotated pseudouridines (Fig. 2C).

Seven Modomics-annotated and DRS-supported Ψ sites, however, failed to pass the threshold in the BACS data (Supplemental Fig. S3A, Supplemental Tables S3&S4). Conversely, six Modomics-annotated Pus1 targets failed to be detected by DRS but were called as pseudouridines by BACS. These results highlight the complementary nature of these approaches for comprehensive characterization.

Two isoacceptors previously annotated to contain Ψ_27_ and corroborated here with BACS did not pass the DRS threshold at U_27_ (Fig. 2C, Supplemental Tables S3&S4). Both isoacceptors did, however, pass the threshold at A_26_ (Supplemental Table S5). This DRS miscall one position proximal to the modified site is likely caused by the loss of Ψ_27_, as there is no annotated modification at position 26 in either isoacceptor. This would be an exception to the predominant observation that miscalls occur coincident with the pseudouridylated base, without affecting signal at neighboring bases (Begik et al. 2021; Shaw et al. 2024; Rübsam et al. 2025).

A recent study performed DRS on tRNA from the fission yeast, *Schizosaccharomyces pombe,* characterizing Pus1-catalyzed modifications using genetic knockouts (Rübsam et al. 2025). We tested whether DRS patterns from budding and fission yeast could clarify a sequence motif for Pus1 modification. Sequence logos of isoacceptors with a U_27_ modified by Pus1 showed a modest preference for a C_25_ in both species. However, there is also a general preference for C_25_ across all cytosolic isoacceptors (Supplemental Fig. S4). Thus, C_25_ is associated with, but not sufficient for modification. This result is consistent with previous work on mRNA sites, which showed that RNA sequence alone is not determinant of modification by Pus1 (Carlile et al. 2019; Grünberg et al. 2023).

### Pus1 does not generally influence the abundance of other tRNA modifications

Deletion of Pus1 led to DRS miscall changes at non-uridine sites including position 26, known to contain methylated guanosines in some tRNAs (Supplemental Table S5, Supplemental Fig. S5). To quantify the extent of other possible changes with the loss of Pus1, we performed total nucleoside liquid chromatography tandem mass spectrometry (LC-MS/MS) on tRNA isolated from wild-type and *pus1*Δ cells (Fig. 2D, Supplemental Table S6). We digested gel-purified tRNA molecules down to mono-ribonucleosides using a two-step enzymatic digestion (Jones et al. 2023a). The nucleosides were then separated using reverse-phase liquid chromatography and analyzed with triple quadrupole mass spectrometry. Here, we quantified twenty-seven different modified nucleoside abundances between strains (Supplemental Table S6).

As expected, *pus1*Δ tRNA had decreased pseudouridine content relative to wild-type. No other modifications showed notable changes in abundance between *pus1*Δ and wild-type (Fig. 2D). Twenty-one non-pseudouridine sites passed the DRS threshold change in *pus1*Δ relative to wild-type (Supplemental Table S5). The lack of changes observed by LC-MS/MS may arise from the fact that any individual site change (i.e. a change in the proportion of a specific isoacceptor whose molecules contain a modification at a specific position), or even changes of the same type of modification across multiple tRNAs, may not result in a quantitatively significant change in total fractional abundance of that modification within the entire tRNA nucleoside pool. Alternatively, some of the changes at certain tRNA positions suggested by DRS could be spurious results arising from sequencing artifacts in certain contexts.

Methylated guanosines, such as m^1^G and m^2,2^G, can induce reverse transcription (RT) stops. We thus leveraged the BACS data from untreated samples of wild-type and *pus1*Δ tRNA to test, at the isoacceptor level, whether such modifications were changed in *pus1*Δ tRNA. We observed poor agreement between DRS signal changes and changes in RT stops between *pus1*Δ and wild-type, indicating the changes observed by DRS are not due to changes in modification abundance (Supplemental Table S7).

### Nanopore DRS and BACS identify both known and unknown Pus7 target sites

We next performed DRS and BACS on cytosolic tRNAs isolated from the *pus7*Δ strain (Fig. 3A&B, Supplemental Fig. S6). Pus7 modifies tRNA, mRNA, 5S rRNA, and U2 snRNA through recognition of specific sequence motifs (Behm-Ansmant et al. 2003; Yang et al. 2005; Decatur and Schnare 2008; Urban et al. 2009; Wu et al. 2011; Schwartz et al. 2014; Carlile et al. 2014; Purchal et al. 2022; Ruan et al. 2026). Pus7 typically localizes to the nucleus but is also found in the cytoplasm under specific stress conditions (Schwartz et al. 2014; Ruan et al. 2026).

**Figure 3.**
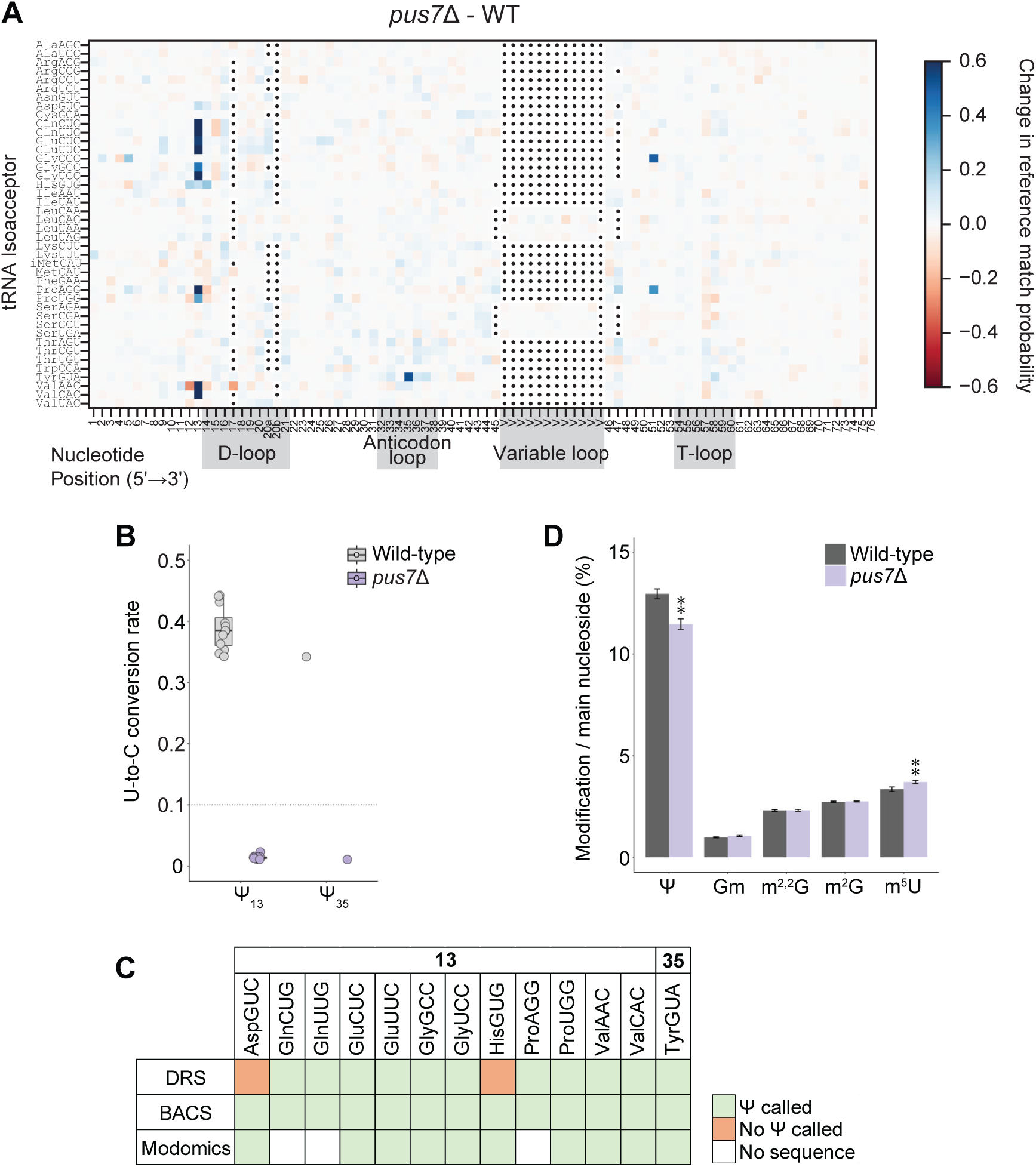
Modification of *pus7*Δ and wild-type *S. cerevisiae* nuclear-encoded tRNA. (A) Subtractive heatmap comparing Nanopore direct RNA sequencing (DRS) reference match probability of *pus7*Δ relative to wild-type. Otherwise as described in Figure 2A. (B) U-to-C conversion rates from 2-bromoacrylamide-assisted cyclization sequencing (BACS) of wild-type and *pus7*Δ cytosolic tRNA. Otherwise as described in Figure 2B. *P* < 0.0001 for all sites shown, comparing wild-type and *pus7Δ* conversion rates (Šídák’s multiple comparisons test). (C) Summary of DRS and BACS-detected, and Modomics-annotated pseudouridine sites catalyzed by Pus7. Otherwise as described in Figure 2C. (D) Total nucleoside LC-MS/MS of tRNA harvested from wild-type and *pus7*Δ strains. Otherwise as described in Figure 2D.

Pus7 is known to modify nine tRNAs at position 13 and one tRNA, tRNA^Tyr(GUA)^, at position 35 (Fig. 1) (Behm-Ansmant et al. 2003; Urban et al. 2009; Wu et al. 2011; Cappannini et al. 2024). DRS predicted 7/9 Modomics-annotated and three previously unannotated Ψ_13_ sites, as well as the single Ψ_35_ on tRNA^Tyr(GUA)^ (Fig. 3A&C, Supplemental Table S2). All Modomics-annotated and the three novel DRS-predicted sites were confirmed with BACS (Fig. 3B&C, Supplemental Table S4).

tRNA^His(GUG)^ was not predicted by DRS to contain Ψ_13_, but was confirmed to contain Ψ_13_ by BACS (Fig. 3B&C, Supplemental Table S4). tRNA^His(GUG)^ did, however, show DRS miscall decreases at adjacent positions A_12_ and A_14_ (Supplemental Table S5). Since there are no annotated modifications at either of these positions, these changes may be the result of the loss of Ψ at position 13. This may represent an additional example of a pseudouridine miscall appearing adjacent to the pseudouridylated base, in contrast to the predominant pattern of pseudouridine miscalls occurring coincident with the modified base (Begik et al. 2021; Shaw et al. 2024; Rübsam et al. 2025; Reinsch and Garcia 2026).

### Pus7 does not generally influence the abundance of other tRNA modifications

Deletion of Pus7 led to changes in DRS signal at eight sites not known to be modified by any PUS (Supplemental Table S5). By total nucleoside LC-MS/MS, *pus7*Δ tRNA showed decreased pseudouridine content relative to wild-type (Fig. 3D). The only other notable change observed was an increase in 5-methyluridine (m^5^U) in *pus7*Δ (11.5%, *P* < 0.0001). m^5^U, catalyzed by Trm2 in budding yeast, is annotated to occur at position 54 in most tRNAs, as measured by mass spectrometry (Nordlund et al. 2000). However, it has not been consistently recorded using Nanopore DRS-based miscall analysis, with variable outcomes between studies (Lucas et al. 2024; White et al. 2024; Shaw et al. 2024; Reinsch and Garcia 2026).

Deletion of Pus7 led to an unexpected decrease in DRS base miscalls at G_51_ in tRNA^Gly(CCC)^ and tRNA^Pro(AGG)^ in our *S. cerevisiae* data, collected using the ONT RNA002 sequencing chemistry (Fig. 3A; Supplemental Table S5). We do not believe this was due to a previously unannotated methylated guanosine, as we did not observe RT stops at G_51_ (Supplemental Table S7).

Notably, a *pus7*Δ-induced change at G_51_ has also been observed in *S. pombe* using ONT RNA004 chemistry in two isoacceptors: tRNA^Asp(GUC)^ and tRNA^Gly(GCC)^ (Rübsam et al. 2025). This may be an effect of structural changes that occur with the loss of Ψ_13_, which has been shown to promote interactions between the D-arm (where Ψ_13_ is located) and T-arm (where G_51_ is located) (Biela et al. 2025). In *S. cerevisiae*, tRNA^Gly(CCC)^, however, is not known to be modified by Pus7 and does not contain a U_13_.

### Nanopore DRS and BACS identify both known and unknown Pus3 target sites

We next sequenced tRNA from *pus3*Δ cells (Fig. 4A&B, Supplemental Fig. S7A). Pus3 modifies tRNAs at positions 38 and 39 (Fig. 1), affecting decoding and frameshifting during protein synthesis (Lecointe et al. 1998, 2002; Han et al. 2015). In budding yeast, loss of Pus3 leads to growth impairment, increased sensitivity to heat, and changes in lipidome structure, protein homeostasis, cell morphology, and stress response (Lecointe et al. 2002; Han et al. 2015; Mülleder et al. 2016; Bruch et al. 2020; Matos et al. 2023). Seven isoacceptors lacking sequences in Modomics contain uridine at position 38 or 39 and have not yet been investigated as potential Pus3 targets.

**Figure 4.**
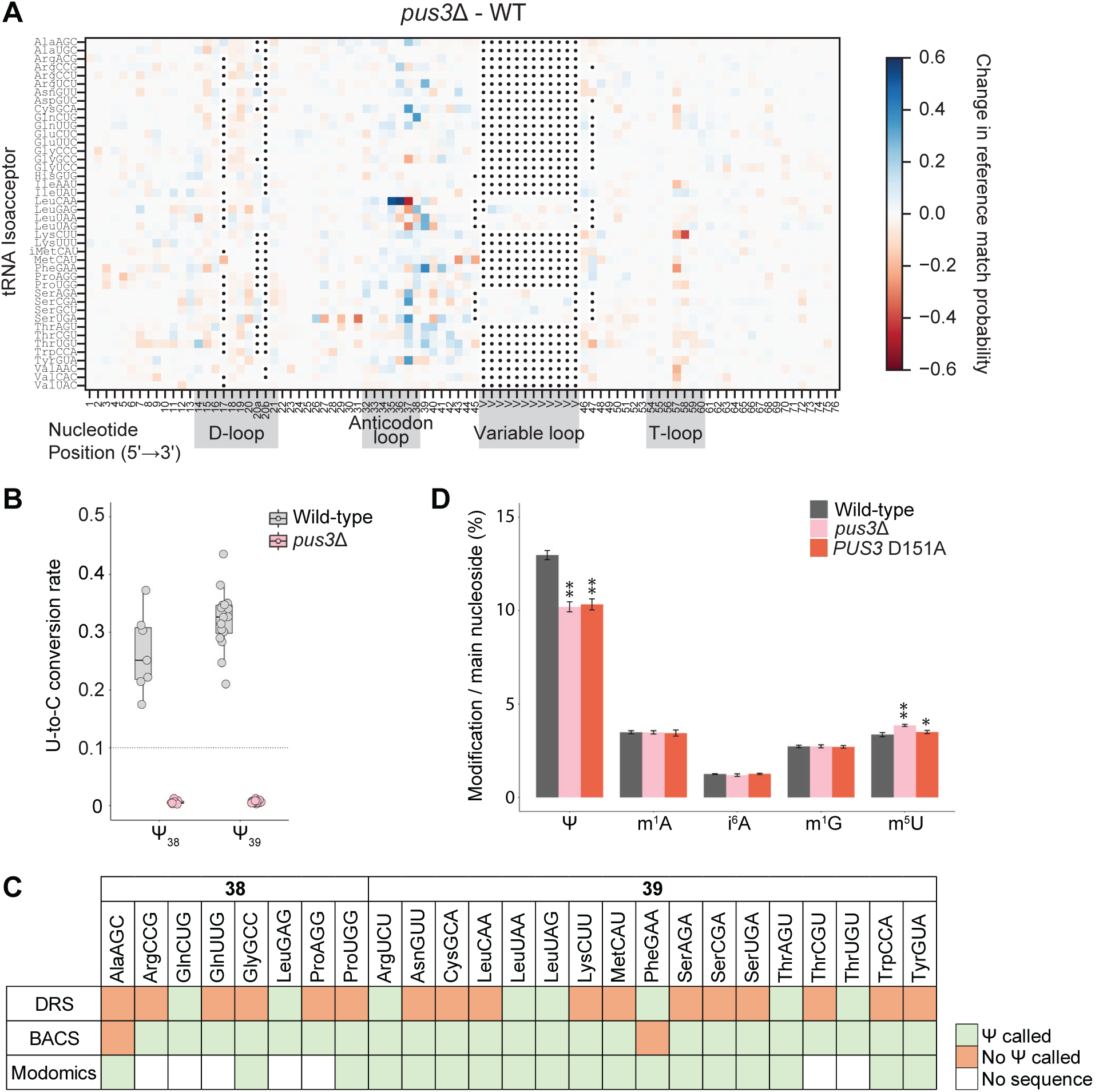
Modification of *pus3*Δ and wild-type *S. cerevisiae* nuclear-encoded tRNA. (A) Subtractive heatmap comparing Nanopore direct RNA sequencing (DRS) reference match probability of *pus3*Δ relative to wild-type. Otherwise as described in Figure 2A. (B) U-to-C conversion rates from 2-bromoacrylamide-assisted cyclization sequencing (BACS) of wild-type and *pus3*Δ cytosolic tRNA. *P* < 0.0001 for all sites shown, comparing wild-type and *pus3*Δ conversion rates (Šídák’s multiple comparisons test). Conversion rates of annotated pseudouridine sites that failed to pass the threshold are shown in Supplemental Figure S3B. Otherwise as described in Figure 2B. (C) Summary of DRS and BACS-detected, and Modomics-annotated pseudouridine sites catalyzed by Pus3. Otherwise as described in Figure 2C. (D) Total nucleoside LC-MS/MS of wild-type, *pus3*Δ, and *PUS3* D151A tRNA. Modification abundances of pseudouridine (Ψ), 1-methyladenosine (m^1^A), N6-isopentenyladenosine (i^6^A), 1-methylguanosine (m^1^G), and 5-methyluridine (m^5^U) are shown. “*” indicates *P* < 0.05, “**” indicates *P* < 0.001 in Dunnett’s multiple comparison test of the mutant values compared to wild-type.

DRS generally did poorly at predicting Modomics-annotated Pus3 sites, while BACS confirmed nearly all annotated sites (Fig. 4C, Supplemental Table S3). Only 5/18 Modomics-annotated sites passed the DRS threshold change in reference match probability, compared to 18/24 for Pus1 and 8/10 for Pus7 (Figs. 2C,3C,&4C, Supplemental Table S3). This was likely due to the weak miscall signals for Pus3-pseudouridylated sites in wild-type DRS reads (Supplemental Fig. S1). Only 8/18 annotated Pus3 sites met the reference match probability cutoff of <0.7 in wild-type, compared to 20/24 for Pus1 and 10/10 for Pus7 (Supplemental Table S1) (Shaw et al. 2024).

Our observations are consistent with previous work in *S. pombe*, which showed a pattern of weaker DRS miscall signals for sites catalyzed by Deg1 (the fission yeast homolog of Pus3) compared to other PUS-modified sites (Rübsam et al. 2025). Miscall signal strength may be influenced by the local modification landscape; positions 38 and 39 are located next to the frequently modified position 37 (see below), and the anticodon that is also frequently modified (positions 34–36) (Supplemental Fig. S5).

DRS did predict three novel Pus3-catalyzed pseudouridine sites, all of which were confirmed with BACS. BACS also identified an additional four novel pseudouridine sites (Fig. 4C, Supplemental Table S3&S4).

Two previously annotated sites failed to be detected by BACS (Fig. 4C). Both of these sites did nevertheless show a significant decrease in U-to-C conversion rate between the wild-type and *pus3*Δ BACS samples (*P* < 0.0001, Supplemental Fig. S3B, Supplemental Table S4). This may be the result of low modification stoichiometry. Across all pseudouridine sites detected by both methods, we did not observe a correlation between DRS change in reference match probability and BACS U-to-C conversion rates (R^2^ = 0.0112, Supplemental Fig. S3C), in accordance with the fact that neither method has been employed here as a quantitative measure of site stoichiometry.

### Loss of Ψ_38/39_ alters DRS signal at position 37 without affecting corresponding modification abundances

Position 37 showed the most occurrences of signal changes of any single tRNA position in *pus3*Δ relative to wild-type (Supplemental Table S2). Recent work in *S. pombe* has also exhibited a high occurrence of signal changes concentrated at position 37 with the deletion of the Pus3-homolog Deg1, demonstrating that this result is not species-specific (Rübsam et al. 2025).

In our DRS data, the strongest changes occurred at sites annotated to contain 1-methylguanosine (m^1^G; catalyzed by Trm5) and N6-isopentenyladenosine (i^6^A; catalyzed by Mod5) (Fig. 1, Supplemental Figure S7B&C, Supplemental Table S5). All isoacceptors annotated to contain i^6^A_37_ also contain Ψ_39_ and showed a decrease in miscalls at position 37 upon deletion of Pus3. Three of the four leucine isoacceptors, all containing a Ψ_38_ or Ψ_39_, showed an increase in miscalls at annotated m^1^G_37_ sites (Supplemental Figure S7B&C).

To measure the influence of Pus3 activity on m^1^G and i^6^A abundance, we performed LC-MS/MS on tRNA purified from wild-type and *pus3*Δ cells, and from cells expressing a Pus3 catalytic mutant, D151A (Lecointe et al. 2002). *PUS3* D151A and *pus3*Δ strains had a comparable decrease in pseudouridine relative to wild-type, demonstrating complete loss of catalytic activity with the D151A mutation (Fig. 4D). *pus3*Δ showed an increase in m^5^U abundance relative to wild-type, similar in magnitude to that seen for *pus7*Δ (15.1%, *P* < 0.0001). i^6^A and m^1^G did not change in abundance between *pus3*Δ, *PUS3* D151A, and wild-type (Fig. 4D; Supplemental Table S6), in contrast to our predictions based on the DRS data. Since all isoacceptors annotated to contain i^6^A exhibited a change in miscall signal at position 37 in *pus3*Δ but no change was observed by LC-MS/MS, we conclude that i^6^A abundance is not affected by the loss of Pus3 activity. This change may instead reflect a change in the effect of i^6^A_37_ on the ionic current during sequencing, in the absence of the Pus3-catalyzed pseudouridine.

m^1^G is annotated to occur at position 37 in a total of seven isoacceptors and at position 9 in fifteen isoacceptors (Supplemental Figure S5) (Cappannini et al. 2024). Thus, a change in m^1^G_37_ abundance in the three leucine isoacceptors may be undetectable by LC-MS/MS due to the high occurrence of unchanged m^1^G at additional sites. To determine the relative abundance of specifically m^1^G_37_ between *pus3*Δ and wild-type cells, we used a primer extension assay to measure m^1^G_37_ in tRNA^Leu(CAA)^, the isoacceptor exhibiting the largest change at position 37 (Supplemental Figure S7B&C). Primer extension assays use reverse transcription stops caused by modifications, such as m^1^G, to measure relative abundances of a modification (Jackman et al. 2003; Swinehart et al. 2013).

We designed an IR800-tagged primer that is complementary to tRNA^Leu(CAA)^, possessing ≥4 nucleotide differences from each of the other three leucine isoacceptors to promote specificity for tRNA^Leu(CAA)^ (Fig. 5A). We observed no significant change in the relative abundance of m^1^G_37_ RT stop products of tRNA^Leu(CAA)^ between wild-type and *pus3*Δ tRNA samples (Fig. 5B; *P* = 0.129). Thus, m^1^G_37_ abundance in tRNA^Leu(CAA)^ does not change upon loss of Pus3, in contrast to what was predicted by DRS. The DRS change at position 37 of tRNA^Leu(CAA)^ may rather reflect a change in how m^1^G_37_ affects the ionic current with the loss of a neighboring Pus3-catalyzed pseudouridine.

**Figure 5.**
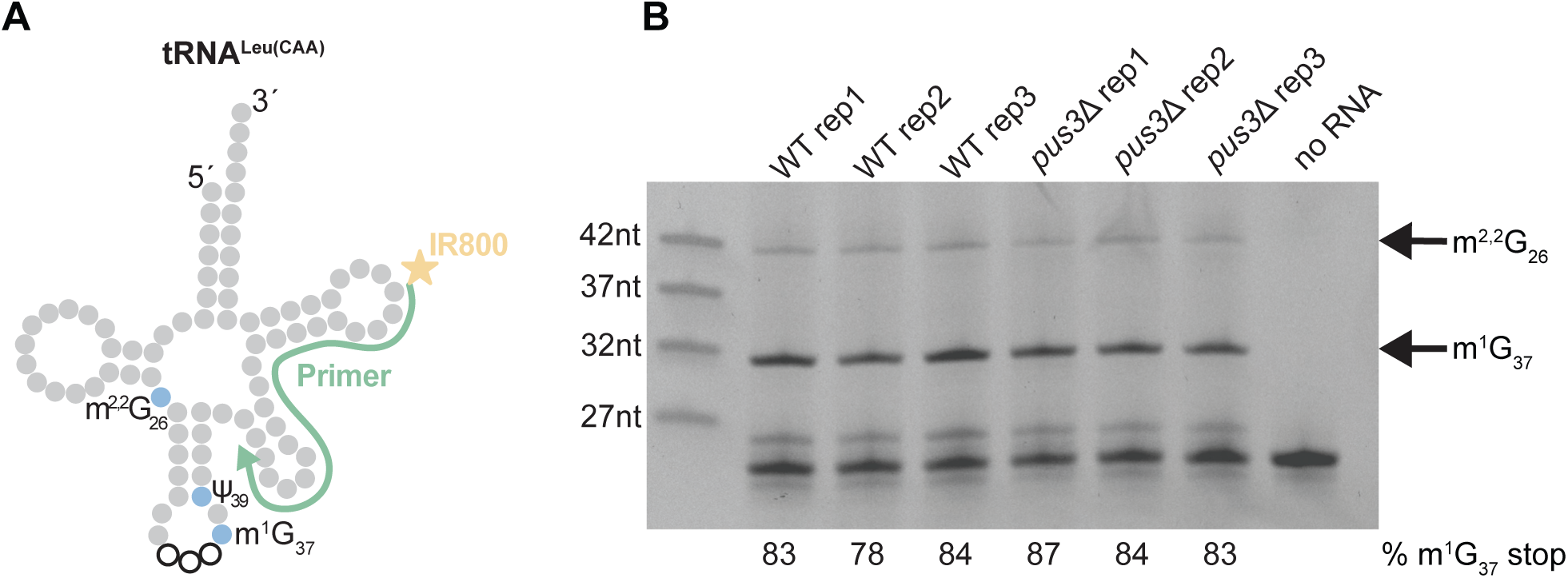
Primer extension analysis of m^1^G_37_ in *S. cerevisiae* tRNA^Leu(CAA)^. (A) Illustration of the IR800-tagged primer that has full complementarity to tRNA^Leu(CAA)^. m^1^G_37_ and m^2,2^G_26_ (modifications causing reverse transcriptase (RT) stops) and Ψ_39_ catalyzed by Pus3 are labeled. (B) tRNA^Leu(CAA)^ primer extension assay with three biological replicates of wild-type and *pus3*Δ tRNA. Gel image shown is the overlay of SYBR gold staining and IR800 signal, aligned using corner slices made in the gel (not shown). Negative control “no RNA" reaction contained no RNA template. Bands indicating RT stops at m^1^G_37_ (28 nt) and m^2,2^G_26_ (39 nt) are labeled with black arrows. Wild-type and *pus3*Δ % stop were compared with a paired one-tailed student’s t-test (*P* = 0.129). The IR800 dye molecule (M.W. = 1163.3 g/mol) causes reverse transcription products to run larger than the ladder oligonucleotides that do not contain this dye. No modifications are known to exist between the 3’ end of the primer and position 39; the faint bands just above the primer-only bands may arise from the secondary structure at the beginning of the anticodon stem interrupting reverse transcription.

We similarly did not observe any RT stop changes between the BACS untreated wild-type and *pus3*Δ samples that corresponded to changes observed by DRS, further suggesting that the DRS changes were not due to changes in the abundance of RT-stop-inducing modifications (Supplemental Table S7).

Overall, these results emphasize the importance of orthogonal methods to test hypothesized modification interdependencies that arise from analysis of DRS data, particularly among closely positioned modifications.

### BACS analysis recapitulates previous annotations of m^1^A_58_

In prior work, we documented the ability of DRS to predict relative changes in m^1^A_58_ in the T-loop (Shaw et al. 2024). BACS treatment can induce conversion of m^1^A to m^6^A, leading to decreased m^1^A-induced mutation rates in BACS treated compared to control samples (Xu et al. 2024). Our wild-type treated samples showed a significant decrease in mutation rate at position A_58_ for all isoacceptors annotated to contain m^1^A_58_ (*P* < 0.0001, Supplemental Fig. S3D, Supplemental Table S8) (Behrens et al. 2021; Cappannini et al. 2024; Shaw et al. 2024). In contrast, all sites annotated to *not* contain m^1^A_58_ did not have statistically significant changes in mutation rate between wild-type treated and untreated samples (*P* > 0.86 for all isoacceptors), with the exception of tRNA^Pro(AGG)^ (*P* = 0.0222).

We leveraged our untreated BACS data to test whether DRS signal changes at annotated m^1^A_58_ sites in *pus3*Δ were reflective of true changes in modification abundance (Fig. 4A, Supplemental Table S5). One isoacceptor, tRNA^Lys(CUU)^, showed a large increase in mutation rate in *pus3*Δ relative to wild-type at A_58_ (112%, *P* < 0.0001) as well as a large DRS signal change (Fig. 4A, Supplemental Tables S5&S8). We have previously documented an increase in m^1^A_58_ on tRNA^Lys(CUU)^ in yeast subjected to nutrient depletion conditions, as measured by DRS (Shaw et al. 2024). Together, these data demonstrate that this particular tRNA is subject to changes in m^1^A_58_ in at least two conditions. tRNA^Phe(GAA)^ showed a more modest increase in mutation rate (15%, *P* < 0.0001), in agreement with the signal change observed by DRS. These results support an increase in m^1^A_58_ abundance with the loss of Pus3 for tRNA^Lys(CUU)^ and tRNA^Phe(GAA)^. None of the other three isoacceptors with changes at A_58_ in DRS, however, showed complementary changes in mutation rate between wild-type and *pus3*Δ samples (Supplemental Table S8). We note also that while *pus1*Δ and *pus7*Δ did not show changes by DRS at position A_58_ compared to wild-type, three isoacceptors from each strain did demonstrate >10% changes in A_58_ mutation rates relative to wild-type as measured by BACS, including tRNA^Phe(GAA)^ in *pus7*Δ (Fig. 2A&3A, Supplemental Table S8).

### Nanopore sequencing and BACS confirm Pus6-dependent pseudouridylation site

Pus6 is annotated to modify only position 31 of the elongator tRNA^Met(CAU)^ (Fig. 1). Both BACS and DRS supported pseudouridylation of U_31_ in tRNA^Met(CAU)^, confirming Pus6 catalysis at this position (Supplemental Fig. S3E&S8, Supplemental Table S4). There were no notable DRS changes at any other tRNA sites, including the 11 previously unannotated isoacceptors, consistent with prior literature (Ansmant et al. 2001).

### BACS identifies novel Ψ_32_ sites

Pus8 is annotated to modify position 32 in eleven tRNAs (Fig. 1) (Behm-Ansmant et al. 2004; Cappannini et al. 2024). The gene *PUS8* (aka *RIB2* in *S. cerevisiae*) encodes a bifunctional pseudouridine synthase and DRAP deaminase enzyme and is an essential gene in the BY budding yeast strain background (Behm-Ansmant et al. 2004). With our wild-type BACS data, we identified five previously unannotated instances of Ψ_32_ (Supplemental Fig. S9, Supplemental Table S4). Without an appropriate mutant control, however, we are unable to confirm that these modifications are catalyzed by Pus8.

### Deletion of mitochondria-specific pseudouridine synthases does not affect cytosolic tRNA modifications detected by DRS

Pus2, Pus5, and Pus9 are encoded in the nuclear genome but localize to mitochondria and have not been observed to localize to the cytoplasm, nor to modify nuclear-encoded tRNAs (Ansmant et al. 2000; Behm-Ansmant et al. 2004, 2007). Pus2 is a paralog of Pus1 and modifies mitochondrial tRNAs at positions 27 and 28 (Behm-Ansmant et al. 2007; Cappannini et al. 2024; Reinsch and Garcia 2026). Pus5 modifies mitochondrial 21S rRNA at U_2819_ and is not known to modify any tRNAs (Ansmant et al. 2000). Pus9 is a paralog of Pus8 and is annotated to modify six mitochondrial isoacceptors at position 32 (Behm-Ansmant et al. 2004). We observed no changes in nuclear-encoded tRNA miscalls with the deletion of Pus2, Pus5, or Pus9 (Supplemental Fig. S10), indicating that these enzymes do not modify cytosolic tRNAs in a manner that was detectable by DRS.

## DISCUSSION

In this work, we identified twenty-one novel pseudouridine sites catalyzed by Pus1, Pus3, Pus7, and Pus8, and created an updated map of all known pseudouridines (totaling 126) across the forty-two nuclear-encoded *S. cerevisiae* tRNAs (Fig. 6, Supplemental Table S3).

**Figure 6.**
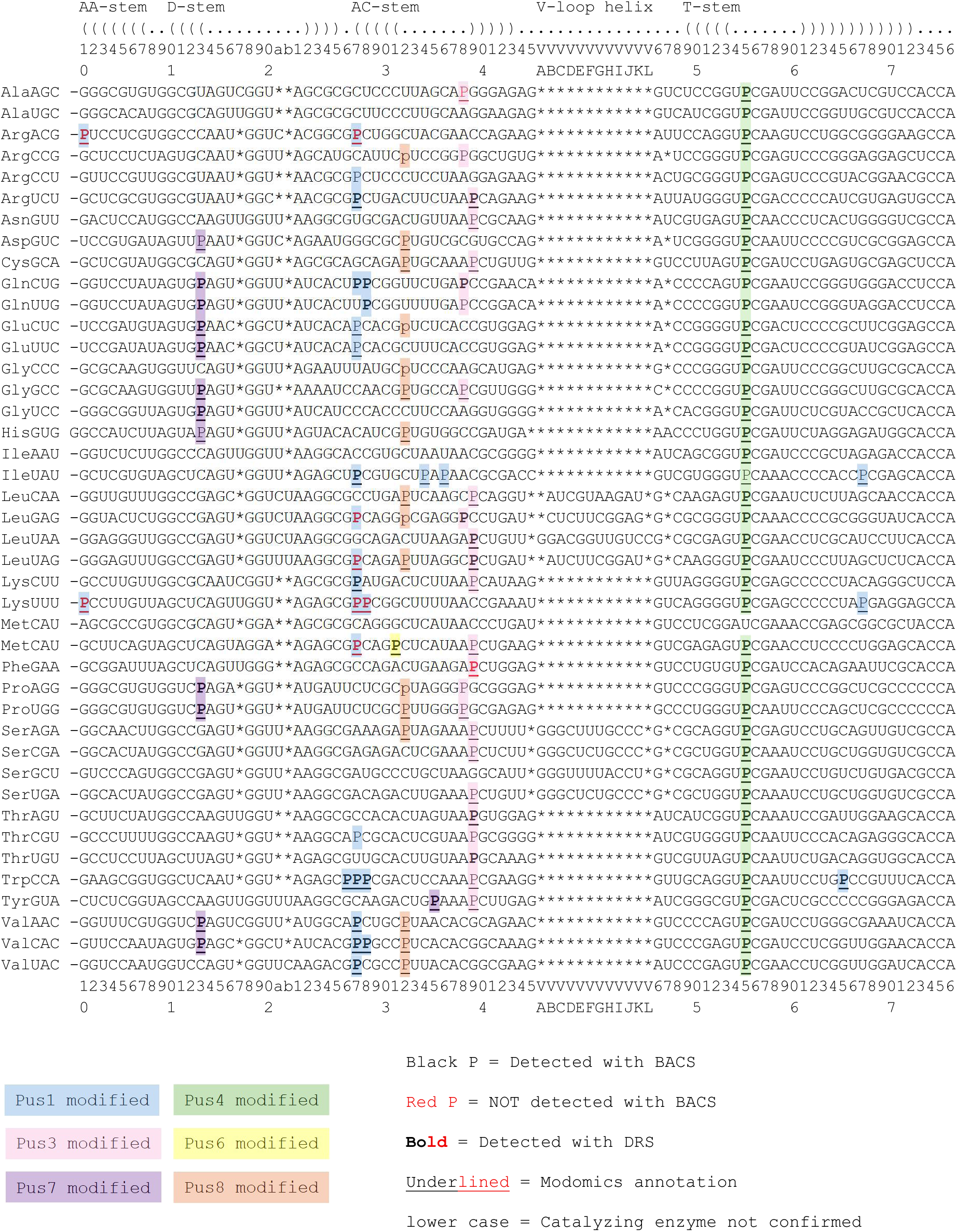
Map of pseudouridines (P) in all 42 *S. cerevisiae* nuclear-encoded tRNAs based on prior annotations combined with 2-bromoacrylamide-assisted cyclization sequencing (BACS) data and direct RNA sequencing (DRS) data from this study and our previous study (Shaw et al. 2024). The enzyme catalyzing P at a given position is denoted by colored boxes. Source of annotation is indicated: Black text (116 sites) = BACS wild-type data from this study. Red text (10 sites) indicates a Modomics and/or DRS pseudouridine annotation that did not pass the threshold to be called as a pseudouridine in the BACS data. **Bold** (82 sites) = DRS PUS deletion/mutant data from this study (Pus1, 3, 6, 7, 8—42 total sites) or in (Shaw et al. 2024) (Pus4—40 total sites). <u>Underlined</u> (95 sites) = Modomics annotation. lower case (5 sites) indicates a pseudouridine annotation for which a PUS protein annotation is not provided in Modomics and could not be resolved by DRS or BACS. All BACS-detected Pus1, Pus3, and Pus7 sites were confirmed to be catalyzed by the corresponding enzyme with BACS PUS deletion data.

Twenty-one sites that have been annotated to contain a pseudouridine in Modomics failed to show evidence of a pseudouridine in the corresponding PUS deletion by DRS. All but one of these sites, however, were successfully detected with BACS. Conversely, nine annotated pseudouridine sites failed to meet the threshold to be detected by BACS but were detected by DRS of PUS deletion strains. Only one Modomics-annotated site failed to be detected by either BACS or DRS (Ψ_38_ in tRNA^Ala(AGC)^). While this may be a result of differences in growth conditions between the annotation source and this study, we note a statistically significant decrease in BACS U-to-C conversion rate was observed for this site between wild-type and *pus3*Δ (*P* < 0.0001, Supplemental Table S4, Supplemental Fig. S3B). This may reflect the presence of a pseudouridine that fell below our BACS threshold to be called as Ψ in this study. Similar results were observed for the nine Pus1- and Pus3-modified pseudouridine sites that were detected by DRS but failed to meet the BACS threshold; these sites nevertheless each showed statistically significant decreases in U-to-C conversion rate in the PUS deletion relative to wild-type (*P* < 0.0001 for each site; Supplemental Table S4, Supplemental Fig. S3A&B). These results emphasize the importance of performing multiple methods to comprehensively capture pseudouridine sites that may otherwise be missed by using one method alone.

Deletion of Pus1, Pus3, and Pus7 in budding yeast led to DRS signal changes both at known PUS-modified sites and at nearby sites (Fig. 2A,3A,4A). Most signal changes were observed to occur at or near positions known to be pseudouridylated. Detection of pseudouridine by DRS base miscalls has been reliably used to identify pseudouridine sites across tRNAs (Lucas et al. 2024; White et al. 2024; Shaw et al. 2024; Reinsch and Garcia 2026). When additional changes in miscall patterns are observed at non-uridine positions with the deletion of a PUS, the causes of these secondary changes are more difficult to establish from DRS data alone. They may reflect 1) signal “crosstalk”, as we described for Pus3 sites neighboring position 37 modifications, 2) modification circuits/interdependencies (Supplemental Fig. 11), or 3) sequencing artifacts. As we have demonstrated here and previously (Shaw et al. 2024; Reinsch and Garcia 2026), orthogonal methods, such as LC-MS/MS and primer extension, can help distinguish between these effects. Additional control molecules, such as synthetic tRNAs lacking specific combinations of modifications, would further clarify the source of these changes.

There are currently widespread efforts to improve Nanopore sequencing accuracy and expand basecalling for specific RNA modifications, including through direct ionic current analysis (Rübsam et al. 2025; Wang et al. 2024; Acera Mateos et al. 2024; Akeson et al. 2025). For these advances to continue to expand the power of DRS in modification detection, it is important for them to address these more challenging sequencing contexts. As this technology continues to develop, orthogonal validation of modification changes predicted from direct RNA sequencing data will continue to represent a powerful approach for extracting the most information from this type of data.

## MATERIALS AND METHODS

### Strain construction

Haploid *S. cerevisiae* BY4741 *pus1*Δ, *pus3*Δ, *pus5*Δ, *pus6*Δ, *pus7*Δ, and *pus9*Δ strains were obtained from the KanMX yeast deletion collection (Giaever et al. 2002). Deletion of the specified PUS-encoding gene was confirmed by Sangar sequencing.

The *PUS3* D151A point mutation was introduced to wild-type BY4741 with a CRISPR plasmid targeting the D151 codon of *PUS3,* derived from pJH2972 (http://www.addgene.org/100956). Oligonucleotides and gBlock were purchased from IDT (Supplemental Table S9). CRISPR plasmid construction was done following the protocol from reference (Anand et al. 2017). Wild-type BY4741 cells were transformed using the lithium acetate method with the CRISPR plasmid targeting the aspartic acid codon (GAC) encoding amino acid 151 of Pus3, along with an 81 bp template containing the alanine codon (GCC) mutation with surrounding homology to *PUS3*.

Transformants were selected on SD–URA agar plates. Introduction of the *PUS3* D151A mutation was confirmed in transformants by Sangar sequencing of the *PUS3* locus. Loss of the CRISPR plasmid was promoted by counterselection on 5-FOA agar plates. Mass spectrometry analysis (IDeA National Resource for Quantitative Proteomics) confirmed wild-type and *PUS3* D151A strains to have comparable levels of the Pus3 protein (Supplemental Fig. S12).

### Yeast growth conditions

Cultures for DRS, LC-MS/MS and BACS were all prepared by the same method. Details of yeast culturing and growth were identical to those described in reference (Shaw et al. 2024). Briefly, yeast were grown at 30°C in YPD to an OD of 0.7–1.0, then pelleted and washed with 1X PBS. Pellets were flash frozen in liquid nitrogen and stored at -80°C.

### Total RNA purification and tRNA isolation

Total RNA was extracted using the hot phenol method and tRNA was purified with 8% denaturing PAGE and UV shadowing as previously described (Shaw et al. 2024). The same methods were used to isolate tRNA for DRS, LC-MS/MS, BACS, and primer extension. A higher quantity of total RNA (∼300-400 µg in 10 uL) was input for gels in preparation of primer extension samples, as primer extension requires higher tRNA input. tRNA sample concentrations were measured by Nanodrop (DRS) or Qubit (LC-MS/MS, BACS, and primer extension).

### Nanopore direct RNA sequencing and data analysis

Samples were sequenced and analyzed following the methods of (Shaw et al. 2024). Briefly, in three biological replicates for each strain (one replicate for each of *pus2*Δ, *pus5*Δ, and *pus9*Δ), 250 ng of total tRNA were ligated to a set of double-stranded splint adapters (Supplemental Table S9) before ligation to ONT sequencing adapters. Samples were sequenced on MinIONs with SQK-RNA002 chemistry, following manufacturer’s instructions. Nanopore current files were basecalled with Guppy v3.0.3, aligned with BWA-MEM to a custom BY4741 tRNA isoacceptor reference modified from (Shaw et al. 2024) (Supplemental Table S10), and filtered with SAMtools. Aligned read counts by isoacceptor for individual replicates are provided in Supplemental Table S11. marginAlign and marginCaller were used to create alignment and error models, and posterior probability calculations, respectively (Thomas et al. 2021). Reference match probability, change in reference match probability, and heatmaps were generated with Python (https://github.com/nanoniki/tRNA-heatmap-generator).

### Threshold change in reference match probability calculation

A threshold for change in reference match probability was calculated by extracting the reference match probability of Modomics-annotated PUS-modified sites for wild-type and the corresponding PUS deletion data for *pus1*Δ, *pus3*Δ, and *pus7*Δ. These three deletions were used as their enzymes corresponded to the largest number of annotated tRNA sites. Change in reference match probability was calculated as *pusX*Δ – WT (Supplemental Table S1). For each of *pus1*Δ, *pus3*Δ, and *pus7*Δ, change in reference match probabilities for annotated Ψ sites with wild-type reference match probabilities below the threshold of 0.7 (Shaw et al. 2024) were averaged and 1 standard deviation was subtracted from this average. These three values — (Average – Standard Deviation) for *pus1*Δ, *pus3*Δ, and *pus7*Δ—were averaged to get the threshold value of 0.1612. All values included in this calculation are given in Supplemental Table S1. Positions with change in reference match probability >0.1612 or <-0.1612 were extracted for further analysis (Supplemental Table S2). Notes on modification annotations were obtained from Modomics RNA sequence and protein annotations (Cappannini et al. 2024).

### Nucleoside Hydrolysis and Quantification via LC-MS/MS

The method for nucleoside quantification was adapted from a previously described method (Jones et al. 2023a). In brief, purified tRNA (200 ng) from *pus1*Δ, *pus3*Δ, *pus7*Δ, *PUS3* D151A, and wild-type budding yeast strains were first hydrolyzed to mononucleotides with 300 U/μg Nuclease P1 (NEB, 100,000 U/mL), 100 mM ammonium acetate, and 100 μM zinc sulfate, in a 10 μL total volume, at 37°C overnight. Samples were then dephosphorylated with 50 U/μg bacterial alkaline phosphatase (BAP, Invitrogen, 150 U/μL), 100 mM ammonium bicarbonate, and 50 mM zinc sulfate, in a 20 μL total volume, at 37°C for 5 hours. Samples were lyophilized and resuspended in 16 μL 40 nM ^15^N_4_-inosine in water.

Samples were analyzed using an Agilent 1290 Infinity II liquid chromatography system coupled to an Agilent 6460 triple quadrupole mass spectrometer. Samples were separated on a Waters Acquity UPLC HSS T3 column (100 Å, 1.8 μm, 1.0 x 100 mm) with an attached guard column (2.1 x 5 mm, 100 Å, 1.8 μm) at 35°C. Liquid chromatography conditions were as previously described (Supplemental Table S12) (Jones et al. 2023a). Samples were run in positive mode with multiple reaction monitoring. The following MS source settings were used: gas temperature 350°C, gas flow 10 L/min, nebulizer 25 psi, sheath gas temperature 350°C, sheath gas flow 11 L/min, and a capillary voltage of 4 kV. Calibration curves of the canonical nucleosides and modified nucleosides were prepared in 40 nM ^15^N_4_-inosine and used to quantify sample nucleoside concentrations. Nucleoside sources, reaction monitoring parameters, and calibration concentrations are given in Supplemental Tables S13-S15. For each nucleoside, modification/main nucleoside % was calculated by taking the ratio of the concentration of that nucleoside to the sum of the concentrations of all nucleosides that originated from the same canonical base (e.g. uridine + all uridine-derived modifications). For each strain, three technical replicates of three biological replicates were prepared and averaged. Statistical significance and adjusted p-values were determined with 2-way ANOVA and Dunnett’s test using the wild-type as a control.

### Primer extension for detection of m^1^G

A custom ladder was designed with short DNA oligonucleotides ordered from IDT (Supplemental Table S9). Oligonucleotides were gel-purified by running 20 µL of 100 µM oligo for each of the four standards on a 12% polyacrylamide gel. Bands were cut out using UV shadowing and eluted overnight in 900µL 0.3 M NaCl at 4°C, then precipitated in 2.2 mL 100% EtOH at -80°C for two hours. Samples were spun at 13,000 rpm for 35 minutes at 4°C, then the supernatant removed and pellets dried. Pellets were resuspended in 20 µL UltraPure distilled water (Invitrogen), and concentrations measured by Nanodrop with the “ssDNA” setting.

Primer extension reactions (Swinehart et al. 2013) were done by annealing 1.25 µg total tRNA in 4 µL water (or 4 µL water alone for the no RNA control) with 1 µL of 1 µM IR800-labeled primer (IDT; Supplemental Table S9) at 95°C for 5 minutes on a heat block, then the samples (still in the block) were moved to room temperature to cool for 1 hour. Extension reactions were done by mixing 5 µL annealing reaction with 1 µL 10X AMV-RT buffer (NEB), 0.8 µL 5 mM dNTPs (NEB), 0.4 µL (4 units) AMV-Reverse Transcriptase (NEB), and 2.8 µL water, incubated at room temperature for 5 minutes, then at 37°C for two hours. Reactions were quenched by adding 10 µL 2X RNA dye (NEB) and stored at -20°C.

Samples were thawed on ice, then denatured at 75°C for 5 minutes. 10 µL of each sample and 50 ng of each ladder oligo were run on a 0.8 mm thick 12% polyacrylamide gel (17 cm wide x 16 cm tall). The gel was stained with SYBR gold (Invitrogen) and visualized on an Amersham Typhoon for IRLong, then reimaged for SYBR gold. Band intensities were quantified in Fiji-ImageJ. The fraction of extension products that terminated at m^1^G_37_ was calculated by dividing the intensity of the m^1^G_37_ band by the summed intensities of the m^1^G_37_ and m^2,2^G_26_ bands.

### 2-bromoacrylamide-assisted cyclization sequencing (BACS)

BACS was performed as previously described (Xu et al. 2024, 2025), omitting RNA fragmentation since the input was purified tRNA. Briefly, 200 ng tRNA was end-healed with T4 PNK (NEB). Prior to ligation, RNA adapter (Supplemental Table S9) was 5’-phosphorylated with T4 PNK (NEB), purified with Zymo Oligo Clean & Concentrator kit, 5’-adenylated with 5’ DNA Adenylation Kit (NEB), and purified with Zymo Oligo Clean & Concentrator kit. End-healed tRNA was then ligated to 5’-adenylated RNA adapter with T4 RNA Ligase 2, truncated KQ (NEB).

Excess RNA adapter was digested with 5’-deadenylase (NEB) and RecJ_f_ (NEB). Samples were then split into control and BACS treatment groups. BACS samples were treated with a solution of 250 mM 2-bromoacrylamide (Enamine) and 625 mM phosphate buffer (pH 8.5). Treated RNA was double-purified with Micro Bio-Spin P-6 Tris Column (Bio-Rad) and Zymo-IC Column (Zymo).

BACS and control samples were annealed with RT primer (Supplemental Table S9), then reverse transcribed with Maxima H-Reverse Transcriptase (Thermo). Excess RT primer was digested with Exo I (NEB). RNA was hydrolyzed with NaOH, then neutralized with HCl. The cDNA was purified with 1.8x volume RNAClean XP beads (Beckman Coulter), ligated to cDNA adapter (Supplemental Table S9) with T4 RNA Ligase I, high concentration (NEB), then purified again with 1.8x volume RNAClean XP beads. Adapted cDNA was amplified with NEBNext Multiplex Oligos for Illumina (96 Unique Dual Index Primer Pairs) and NEBNext Ultra II Q5 Master Mix for 12 cycles. Amplified cDNA was purified with 1.8x volume Mag-Bind TotalPure NGS beads (Omega Bio-Tek) and sequenced with NextSeq 2000 (60-bp paired end reads).

### BACS data analysis

Read processing was performed as previously described (Xu et al. 2024, 2025). In brief, raw reads were processed with Cutadapt (v5.2) for removal of adapters, and low quality and short reads (-q 20 -m18). Unique molecular identifiers (UMIs) were extracted with UMI-tools extract (v1.1.6) and paired end reads merged with fastp (v1.3.6) with overlap length requirement of 10, overlap difference percent limit of 20, and adapter trimming disabled. Merged reads were then aligned with Bowtie2 (v2.2.5) to the tRNA reference (Supplemental Table S16) with parameters -L 16 -N 1 -mp 4. Aligned reads were sorted and filtered with SAMtools (v1.22.2) to keep reads with mapping quality scores ≥1. Reads were then deduplicated with UMI-tools dedup (v1.1.6). Mutations were counted with SAMtools mpileup (v1.22.2) and cpup (v.0.1.0; https://github.com/y9c/cpup/). Overall read counts and aligned isoacceptor read counts are given in Supplemental Tables S17 and S18, respectively.

Pseudouridine sites were called using U-to-C conversion rates, where conversion rate = C/(C+T) (Supplemental Table S4). Sites with BACS conversion rate ≥0.1, control conversion rate ≤0.01, and control T-to-A/G conversion rates ≤0.1, were called as pseudouridine. Site A_58_ mutation rates were calculated as the ratio of non-A calls at A_58_ to the total read count at that site (C+T+G+gap+deletion+insertion+skip)/(A+C+T+G+gap+deletion+insertion+skip) (Supplemental Table S8). The reverse transcription (RT) stop at a position n was calculated as (read depth)_n_/(read depth)_n+1_. Sites with RT stop values <0.9 are given in Supplemental Table S7. Statistical significance and adjusted p-values for U-to-C conversion rates, mutation rates, and RT stops were determined with 2-way ANOVA and Šídák’s test or Dunnett’s test using the wild-type as a control.

## ACKNOWLEDGEMENTS

We thank Mark Akeson (UC Santa Cruz) for early discussions about the project. We thank Olivia Roumaya, Catherine Edgington, and Jane Jackman (The Ohio State University) for their guidance on the primer extension protocol. We thank Jeff Bishop (Genomics and Cell Characterization Core Facility, University of Oregon) for guidance on Illumina sequencing. We thank Alice Barkan and Julia Widom (University of Oregon) for their comments on the manuscript. Funding for the IDeA National Resource for Quantitative Proteomics was provided through a grant from the National Institute of General Medical Sciences (R24GM137786).

## DATA AVAILABILITY

The direct RNA sequencing and 2-bromoacrylamide-assisted cyclization sequencing data are publicly accessible at the European Nucleotide Archive https://www.ebi.ac.uk/ena (Accession number PRJEB112371).

