## Supplemental Figures for "Pseudouridylation landscape across 42 nuclear-encoded *S. cerevisiae* tRNAs"

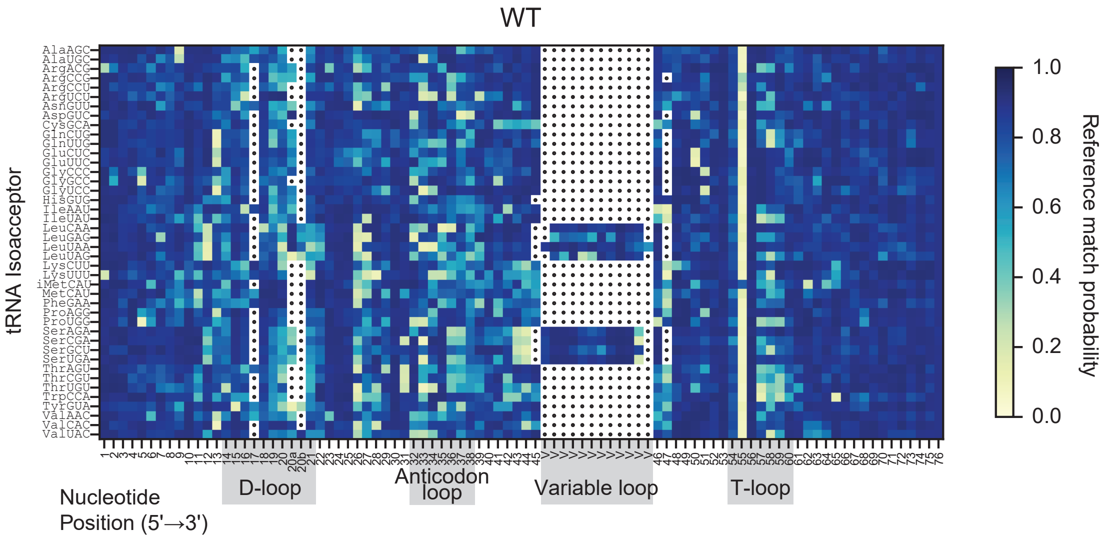


**Figure S1**. Heatmap of Nanopore DRS reference match probabilities of wild-type budding yeast nuclear-encoded tRNA. Reference match probability measures the probability that the DRS-called base matches the canonical base in the reference sequence. A low reference match probability (yellow) is suggestive of a chemical base modification. Data from (Shaw et al. 2024).

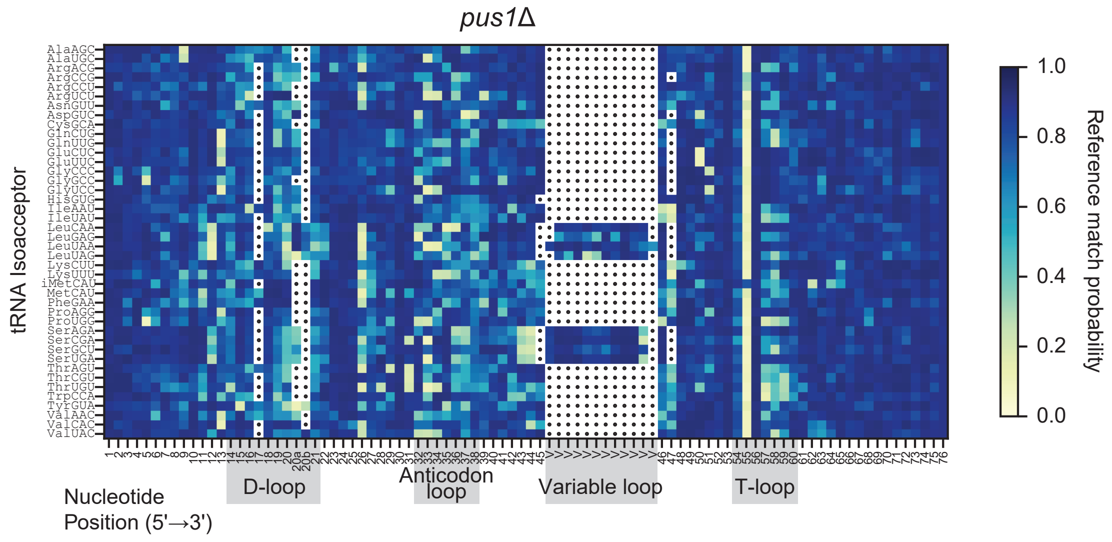


**Figure S2.** Heatmap of Nanopore DRS reference match probabilities of budding yeast nuclear-encoded tRNA from a *pus1*Δ strain. Reference match probability measures the probability that the DRS-called base matches the canonical base in the reference sequence. A low reference match probability (yellow) is suggestive of a chemical base modification.

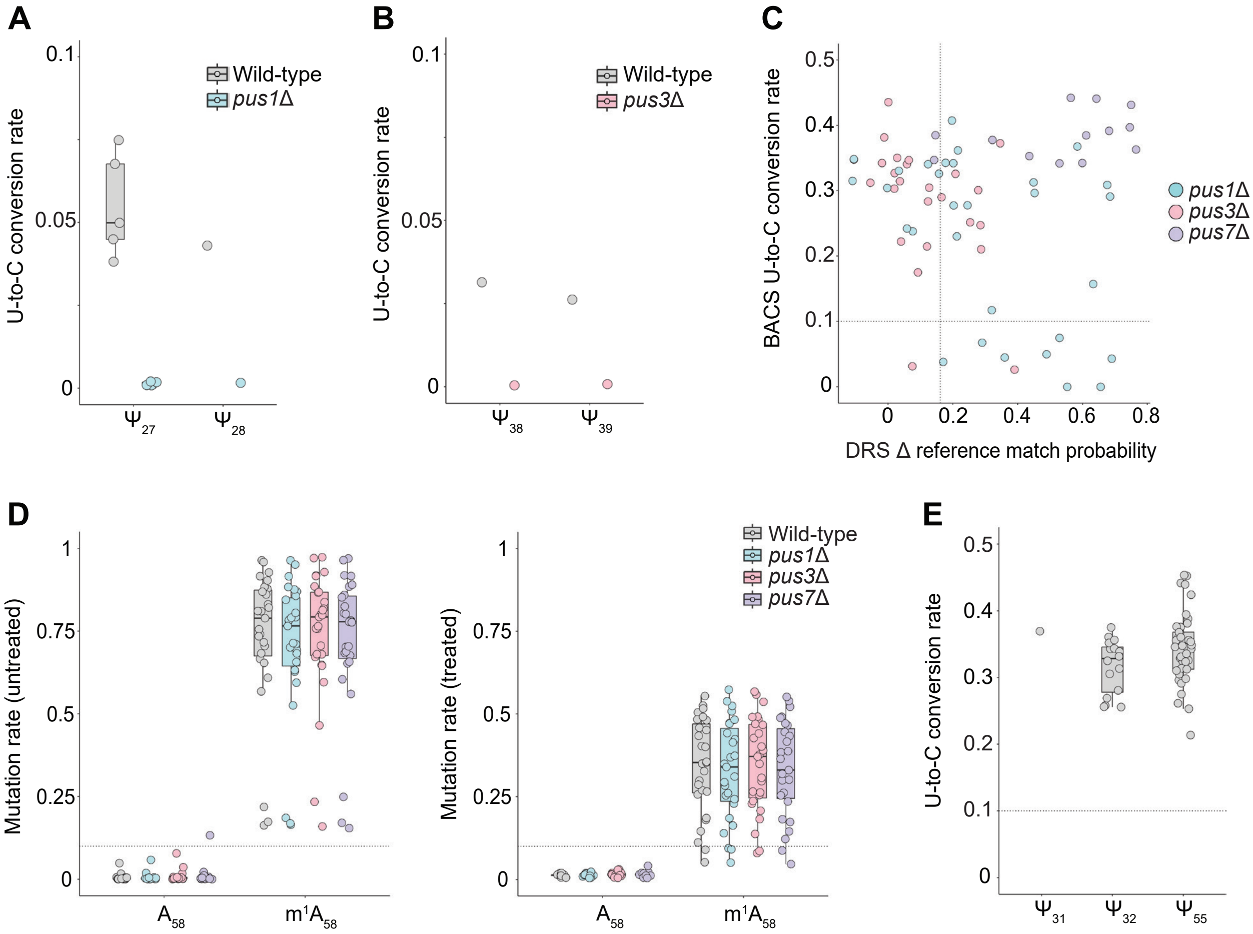


**Figure S3**. 2-bromoacrylamide-assisted cyclization sequencing (BACS) of *S. cerevisiae* nuclear-encoded tRNA. (A) Sites annotated to contain a pseudouridine catalyzed by Pus1 in Modomics and/or DRS data that failed to pass the BACS conversion rate threshold of 0.1. These sites did, however, show decreased conversion in *pus1*Δ, relative to wild-type ($P <$ 0.0001). Three of the included $\Psi_{27}$ sites share a CG**U**CA motif that is not present for any sites that were detected by BACS. Conversion rates for position 1 annotations were zero. Previous BACS studies with human samples have not recorded any pseudouridine at a terminal position (Xu et al. 2024, 2025). Thus, this method may not be able to detect Ψ occurring at the terminal position of a molecule. An additional four currently unannotated Pus1 sites were also implicated in our BACS data—$tRNA^{Asn(GUU)}$, $tRNA^{Gln(UUG)}$, and $tRNA^{Thr(UGU)}$ at position 27 and $tRNA^{Ser(GCU)}$ at position 28 ($P <$ 0.0001)—though these sites failed to meet the threshold cutoff to be called as pseudouridines (Supplemental Table S4). (B) Sites annotated in Modomics to contain a pseudouridine catalyzed by Pus3 that failed to pass the BACS conversion rate threshold of 0.1. These sites did, however, show lower conversion rates in *pus3*Δ, relative to wild-type ($P <$ 0.0001, Supplemental Table S4). (C) DRS change in reference match probability between the indicated PUS deletion and wild-type compared to BACS treated wild-type U-to-C conversion rate. $R^2$ = 0.0112. Each data point is a site at a position targeted by either Pus1, Pus3, or Pus7 that meets at least one of the following criteria: (1) Modomics pseudouridine annotation, (2) Pseudouridine predicted by DRS, or (3) Pseudouridine detected by BACS. Dotted lines denote thresholds for calling Ψ in DRS (0.1612) and BACS (0.1). (D)

Mutation rates at position $A_{58}$ of BACS untreated (left) and treated (right) samples for wild-type, *pus1*Δ, *pus3*Δ, and *pus7*Δ. A decreased mutation rate in treated samples compared to untreated indicates $m^1A_{58}$ modification. Mutation rate calculation is described in **Materials and Methods**. Dashed line marks mutation rate of 0.1. Sites are separated by those annotated to contain $m^1A_{58}$ and those that are not ($A_{58}$) based on previous work (Behrens et al. 2021; Cappannini et al. 2024; Shaw et al. 2024). (E) Pseudouridine sites at positions modified by Pus6 ($\Psi_{31}$), Pus8 ($\Psi_{32}$), and Pus4 ($\Psi_{55}$) detected in wild-type BACS data. (A-E) Each data point from BACS datasets represents a single site on a single isoacceptor (e.g. $\Psi_{39}$ on $tRNA^{Leu(CAA)}$), averaged across three biological replicates. Conversion rate in cDNA reads was calculated as (C/(T+C)), where C is cytosine and T is thymine.

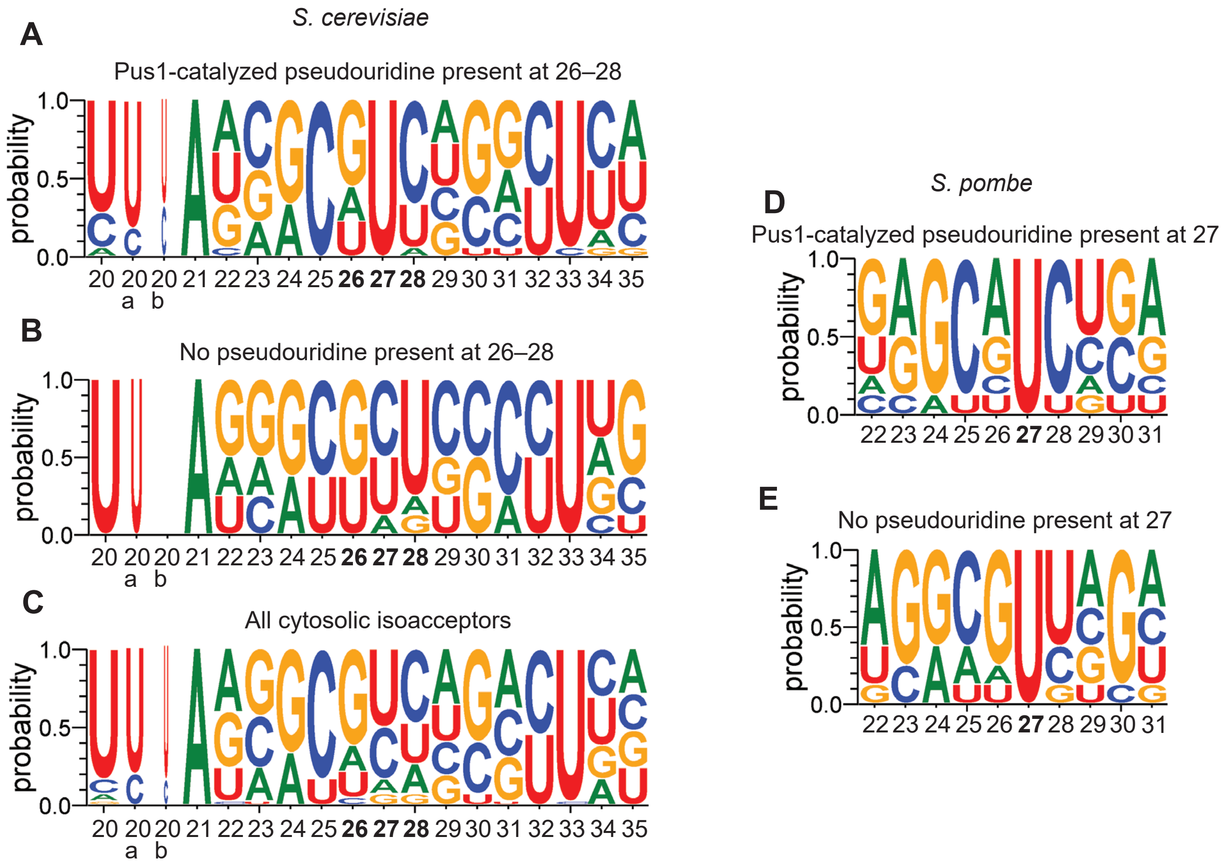


**Figure S4**. Sequence logos of *S. cerevisiae* and *S. pombe* (Rübsam et al. 2025) nuclear-encoded tRNAs for positions surrounding $U_{27}$ based on Pus1 modification status. (A) Sequence logo of the eighteen *S. cerevisiae* isoacceptors containing a pseudouridine at positions 26-28 based on Modomics annotations (Cappannini et al. 2024), and DRS and BACS data from this study. (B) Sequence logo of the eight *S. cerevisiae* isoacceptors containing a $U_{26\text{-}28}$ that are not known to contain a pseudouridine at any of these positions based on Modomics annotations, and BACS and DRS data from this study. (C) Sequence logo of all forty-two *S. cerevisiae* cytosolic isoacceptors. (D) Sequence logo of the eight *S. pombe* isoacceptors containing a Pus1-confirmed pseudouridine at position 27 based on (Rübsam et al. 2025) (E) Sequence logo of the thirteen *S. pombe* isoacceptors containing a $U_{27}$ that is not known to be pseudouridylated based on (Rübsam et al. 2025). Five isoacceptors, $tRNA^{Ala(CGC)}$, $tRNA^{Gln(CUG)}$, $tRNA^{Ile(AAU)}$, $tRNA^{Trp(CCA)}$, and $tRNA^{Val(AAC)}$, contain a $U_{27}$ and showed DRS evidence of $\Psi_{27}$ but were not confirmed Pus1 targets (Rübsam et al. 2025), and thus were excluded from this analysis. Logos generated with WebLogo3 (Crooks et al. 2004).

```
   AA-stem  D-stem             AC-stem             V-loop helix   T-stem
   (((((((..((((..........)))).(((((.......)))))..................(((((.......))))))))))))....
   0        1         2          3         4      ABCDEFGHIJKL    5         6         7
   12345678901234567890ab12345678901234567890123456789012345VVVVVVVVVVVV6789012345678901234567890123456
A -GGGCGUGUKGCGUAGDCGGD**AGCGCRCUCCCUUIGCOPGGGAGAG************GDCUCCGGTPCGAUUCCGGACUCGUCCACCA
A -GGGCACAUGGCGCAGUUGGU**AGCGCGCUUCCCUUGCAAGGAAGAG************GUCAUCGGUUCGAUUCCGGUUGCGUCCACCA
R -PUCCUCGUKLCCCAAD*GGDC*ACGGCRPCUGGCUICGAACCAGAAG************ADU?CAGGTPCAѣGUCCUGGCGGGGAAGCCA
R -GCUCCUCUAGUGCAAU*GGUU*AGCAUGCAUUCUUCCGGUGGCUGUG************A*UCCGGGUUCGAGUCCCGGGAGGAGCUCCA
R -GUUCCGUUGGCGUAAU*GGU**AACGCGUCUCCCUCCUAAGGAGAAG************ACUGCGGGUUCGAGUCCCGUACGGAACGCCA
R -GCUCGCGUKLCGUAAD*GGC**AACGCRPCUGACU1CU6APCAGAAG************ADUAUGGGTPCGѣCCCCCAUCGUGAGUGCCA
N -GACUCCAUGLCCAAGDDGGDD*AAGGCRUGCGACUGUU6APCGCAAG************AD?GUGAGTPCAѣCCCUCACUGGGGUCGCCA
D -UCCGUGAUAGUUPAAD*GGDC*AGAAUGGGCGCPUGUCKCGUGCCAG************A*U?GGGGTPCAAUUCCCCGUCGCGGAGCCA
C -GCUCGUAUKGCGCAGD*GGD**AGCGCAGCAGAPUGCAЧAPCUGUUG************7D?CUUAGTPCGѣUCCUGAGUGCGAGCUCCA
Q -GGUCCUAUAGUGUAGU*GGUU*AUCACUUUCGGUUCUGAUCCGAACA************A*CCCCAGUUCGAAUCCGGGUGGGACCUCCA
Q -GGUCCUAUAGUGUAGU*GGUU*AUCACUUUCGGUUUUGAUCCGGACA************A*CCCCGGUUCGAAUCCGGGUAGGACCUCCA
E -UCCGAUGUAGUGPAAC*GGCU*AUCACAPCACGUUCUCACCGUGGAG************A*C?GGGGUPCGACUCCCCGCUUCGGAGCCA
E -UCCGAUAUAGUGPAAC*GGCD*AUCACAPCACGCU3UCACCGUGGAG************A*C?GGGGTPCGACUCCCCGUAUCGGAGCCA
G -GCGCAAGUKGUUCAGU*GGUU*AGAAUUUAUGCUUCCCAAGCAUGAG************G*CCCGGGUUCGAUUCCCGGCUUGCGCACCA
G -GCGBAAGUKGUUPAGD*GGD**AAAAUCCAACGPUGCCAPCGUUGGG************C*C?CCGGTPCGAUUCCGGGCUUGCGCACCA
G -GGGCGGUUAGUGPAGD*GGDD*AUCAUCCCACCCUNCCAAGGUGGGG************A*CACGGGTPCGAUUCUCGUACCGCUCACCA
H GGCCмUCUUAGUAPAGD*#GDD*AGUACACAUCGPUGUGKCCGAUGA************AAC?CUGGTPCGAUUCUAGGAGAUGGCACCA
I -GGUCUCUUKLCCCAGDDGGDD*AAGGCACCGUGCUIAU6ACGCGGGG************AD?AGCGGTPCGѣUCCCGCUAGAGACCACCA
I -GCUCGUGUALCUCAGD*GGDD*AGAGCUPCGUGCUPAP6ACGCGACC************7D?GUGGGTPCAѣACCCCACCPCGAGCACCA
L -GGUUGUUUGLCMGAGC*#GDCDAAGGCRCCUGAPU?AAKCPCAGGU**AUCGUAAGAU*G*?AAGAGTPCGAAUCUCUUAGCAACCACCA
L -GGUACUCUGGCCGAGU*GGUCUAAGGCGUCAGGUCGAGKUCCUGAU**CUCUUCGGAG*G*CGCGGGUUCAAACCCCGCGGGUAUCACCA
L -GGAGGGUUGLCMGAGD*#GDCDAAGGCRGCAGABUNAAKAPCUGUU*GGACGGUUGUCCG*?GCGAGTPCGѣACCUCGCAUCCUUCACCA
L -GGGAGUUUGLCMGAGD*#GDDDAAGGCRPCAGAPUUAGKCPCUGAU**AUCUUCGGAU*G*?AAGGGTPCGѣAUCCCUUAGCUCUCACCA
K -GCCUUGUUKLCGCAADCGGD**AGCGCRPAUGACUCUU6APCAUAAG************7UUAGGGGTPCGѣGCCCCCUACAGGGCUCCA
K -PCCUUGUUALCUCAGDDGGD**AGAGCRPPCGGCU3UU6ACCGAAAU************7D?AGGGGTPCGѣGCCCCCUAPGAGGAGCCA
M -AGCGCCGUKLCGCAGD*GGA**AGCGCRCAGGGCUCAU6ACCCUGAU************7D??UCGGAUCGѣAACCGŋGCGGCGCUACCA
M -GCUUCAGUALCUCAGDAGGA**AGAGCRPCAGPCUCAU6APCUGAAG************7D?GAGAGTPCGѣACCUCCCCUGGAGCACCA
F -GCGGAUUUALCUCAGDDGGG**AGAGCRCCAGABU#AAYAP?UGGAG************7UC?UGUGTPCGѣUCCACAGAAUUCGCACCA
P -GGGCGUGUGGUCUAGA*GGU**AUGAUUCUCGCUUAGGGUGCGGGAG************GUCCCGGGUUCGAGUCCCGGCUCGCCCCCCA
P -GGGBGUGUKGUCPAGD*GGD**AUGAUUCUCGCPUNGGKPGCGAGAG************7CCCUGGGTPCAѣUUCCCAGCUCGCCCCCCA
S -GGCAACUUGGCMGAGD*#GDD*AAGGCRAAAGAPUIGAЧAPCUUUJ*GGGCUUUGCCC*G*?GCAGGTPCGAGUCCUGCAGUUGUCGCCA
S -GGCACUAUGGCMGAGD*#GDD*AAGGCRAGAGAЩUCGAЧAPCUCUJ*GGGCUCUGCCC*G*?GCUGGTPCAAAUCCUGCUGGUGUCGCCA
S -GUCCCAGUGGCMGAGD*#GDD*AAGGCRAUGCCCUGCU6AGGCAUJ*GGGUUUUACCU*G*?GCAGGTPCGѣAUCCUGUCUGUGACGCCA
S -GGCACUAUGGCMGAGD*#GDD*AAGGCRACAGAЩU&GAЧAPCUGUJ*GGGCUCUGCCC*G*?GCUGGTPCAAAUCCUGCUGGUGUCGCCA
T -GCUUCUAUKLCCAAGDDGGD**AAGGCRCCACAЩUIGU6APGUGGAG************AD?AUCGGTPCAѣAUCCGAUUGGAAGCACCA
T -GCCCUUUUKGCCAAGU*GGU**AAGGCAUCGCACUCGUAAUGCGGGG************AUCGUGGGUUCAAUUCCCACAGAGGGCACCA
T -GCCUCCUUAGCUUAGU*GGU**AGAGCGUUGCACUUGUAAUGCAAAG************GUCGUUAGUUCAAUUCUGACAGGUGGCACCA
W -GAAGCGGUKLCUCAAD*#GD**AGAGCPPPCGABUBCAAAPCGAAGG************7DUGCAGGTPCAѣUUCCUGPCCGUUUCACCA
Y -CUCUCGGUALCCAAGDD#GDDDAAGGCRCAAGACUGPAЧAPCUUGAG************AD?GGGCGTPCGѣCUCGCCCCCGGGAGACCA
V -GGUUUCGUKGUCPAGDCGGDD*AUGGCAPCUGCPUIACACGCAGAAC************7DC?CCAGTPCGѣUCCUGGGCGAAAUCACCA
V -GUUCCAAUALUGPAGC*GGCD*AUCACLPPGCCPUCACACGGCAAAG************7DC?CGAGTPCGѣUCCUCGGUUGGAACACCA
V -GGUCCAAUKLUCCAGD*GGDDCAAGACRPCGCCPU&ACACGGCGAAG************ADC?CGAGTPCGѣACCUCGGUUGGAUCACCA
   12345678901234567890ab12345678901234567890123456789012345VVVVVVVVVVVV6789012345678901234567890123456
   0        1         2          3         4      ABCDEFGHIJKL    5         6         7
```

X = No sequence available in Modomics

X = Modomics sequence annotation

X = Modomics protein annotation only

**Figure S5.** Map of all Modomics-annotated modifications in yeast cytosolic tRNAs, superimposed on a structural alignment from (Shaw et al. 2024). One-letter codes of all annotated Modomics modifications are shown in each indicated position with key listed below (Cappannini et al. 2024). Yellow-highlighted bases are annotated to be modified; Blue-highlighted bases are named in the annotation of the catalyzing enzyme but not in the tRNA sequence annotation; Gray-highlighted isoacceptors do not have sequences available on

Modomics. Anticodon bases are shown in bold. tRNA structural domains are denoted above the sequence positions.

K – m1G; 1-methylguanosine
D – D; dihydrouridine
R - m2,2G; N2,N2-dimethylguanosine
I – I; inosine
O – m1I; 1-methylinosine
P – Ψ; pseudouridine
T – m5U; 5-methyluridine
? – m5C; 5-methylcytidine
Ѣ – m1A; 1-methyladenosine
L – m2G; N2-methylguanosine
1 - mcm5U; 5-methoxycarbonylmethyluridine
6 – t6A; N6-threonylcarbamoyladenosine
Ч – i6A; N6-isopentenyladenosine
7 – m7G; 7-methylguanosine
3 - mcm5s2U; 5-methoxycarbonylmethyl-2-thiouridine
B – Cm; 2'-O-methylcytidine
N – xU; unidentified uridine modification
ʍ - Am; 2'-O-methyladenosine
### - Gm; 2'-O-methylguanosine
M – ac4C;  N4-acetylcytidine
ŋ - Ar(p); 2'-O-ribosyladenosine (phosphate)
Y – yW; wybutosine
Щ – m3C; 3-methylcytidine
J – Um; 2'-O-methyluridine

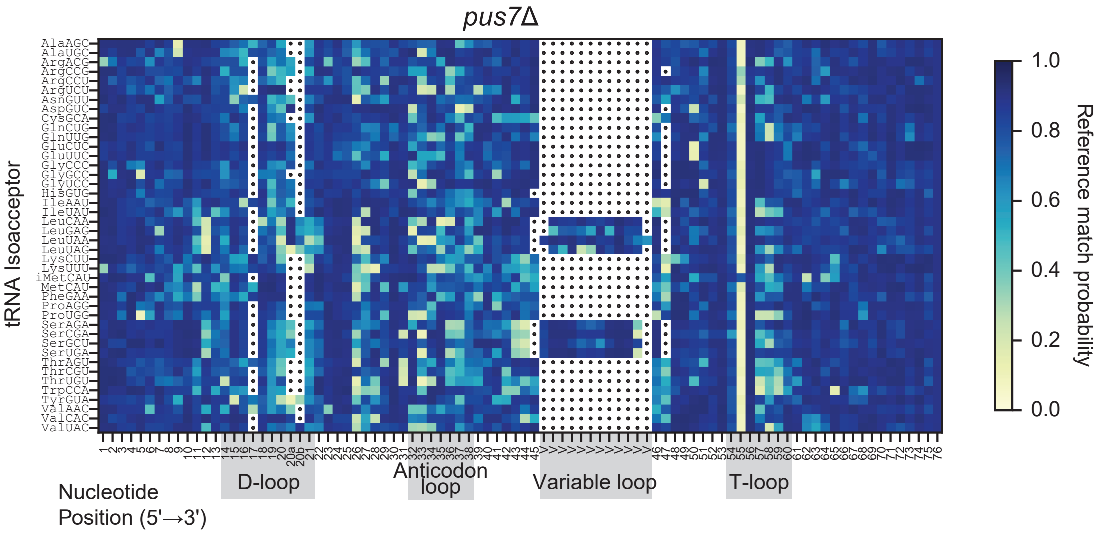


**Figure S6.** Heatmap of Nanopore DRS reference match probabilities of budding yeast nuclear-encoded tRNA from a *pus7*Δ strain. Reference match probability measures the probability that the DRS-called base matches the canonical base in the reference sequence. A low reference match probability (yellow) is suggestive of a chemical base modification.

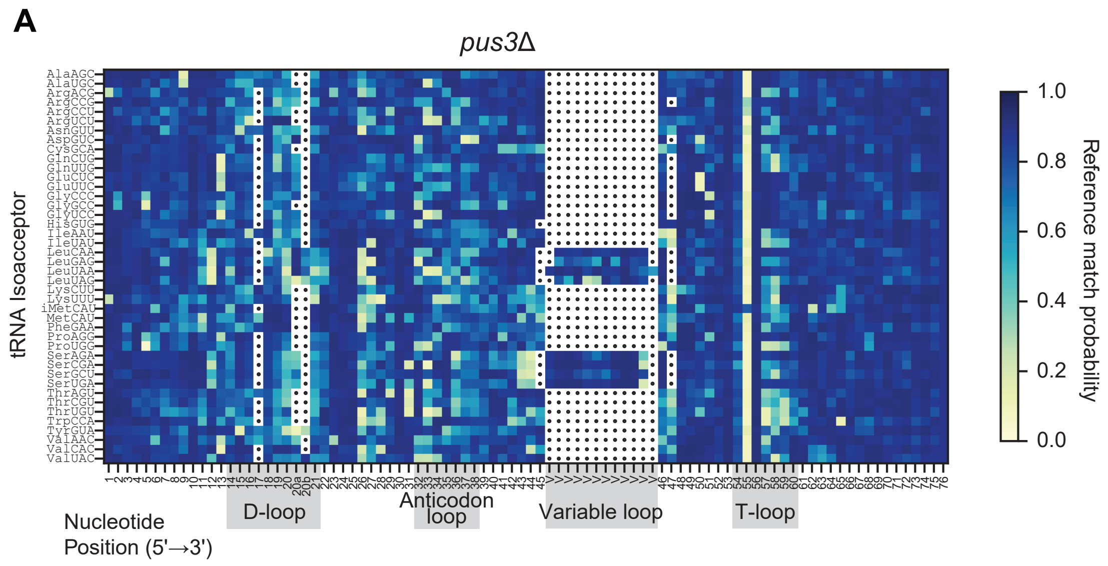


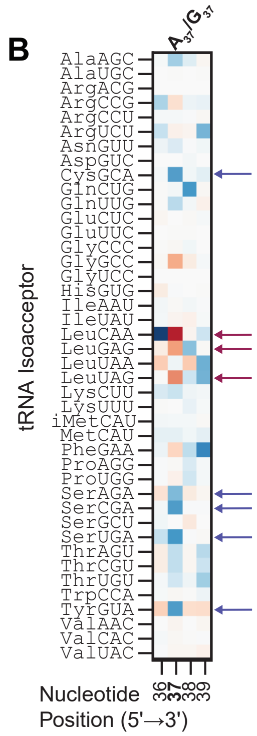


C

| Isoacceptor | Δ reference match probability | Base | Annotation | Pus3 Ψ |
|---|---|---|---|---|
| AlaAGC | 0.203 | A | $m^1I$ | Ψ38 |
| CysGCA | 0.33 | A | $i^6A$ | Ψ39 |
| GlnUUG | 0.164 | A | - | Ψ38 |
| GlyGCC | -0.221 | A | - | Ψ38 |
| LeuCAA | -0.468 | G | $m^1G$ | Ψ39 |
| LeuGAG | -0.252 | G | $m^1G$ | Ψ38 |
| LeuUAG | -0.289 | G | $m^1G$ | Ψ39 |
| SerAGA | 0.263 | A | $i^6A$ | Ψ39 |
| SerCGA | 0.337 | A | $i^6A$ | Ψ39 |
| SerUGA | 0.345 | A | $i^6A$ | Ψ39 |
| TyrGUA | 0.336 | A | $i^6A$ | Ψ39 |

**Figure S7**. Nanopore direct RNA sequencing of budding yeast nuclear-encoded tRNA from a *pus3*Δ strain. (A) Heatmap of Nanopore DRS reference match probabilities of *pus3*Δ tRNA. Reference match probability measures the probability that the DRS-called base matches the canonical base in the reference sequence. A low reference match probability (yellow) is suggestive of a chemical base modification. (B) Enlarged subtractive heatmap of reference match probability for *pus3*Δ minus wild-type tRNA for positions 36-39. Blue arrows indicate threshold-passing positions annotated to contain $i^6A_{37}$. Red arrows indicate threshold-passing positions annotated to contain $m^1G_{37}$. Heatmap scale as in Fig. 4A. See Fig. 4A for full heatmap. (C) Table showing isoacceptors with a change in reference match probability at position 37 that passed the empirically determined threshold value (Materials and Methods), with canonical base, Modomics-annotated modification, and Pus3-catalyzed modification of the specified isoacceptor. “-“ indicates no annotated modification in Modomics.

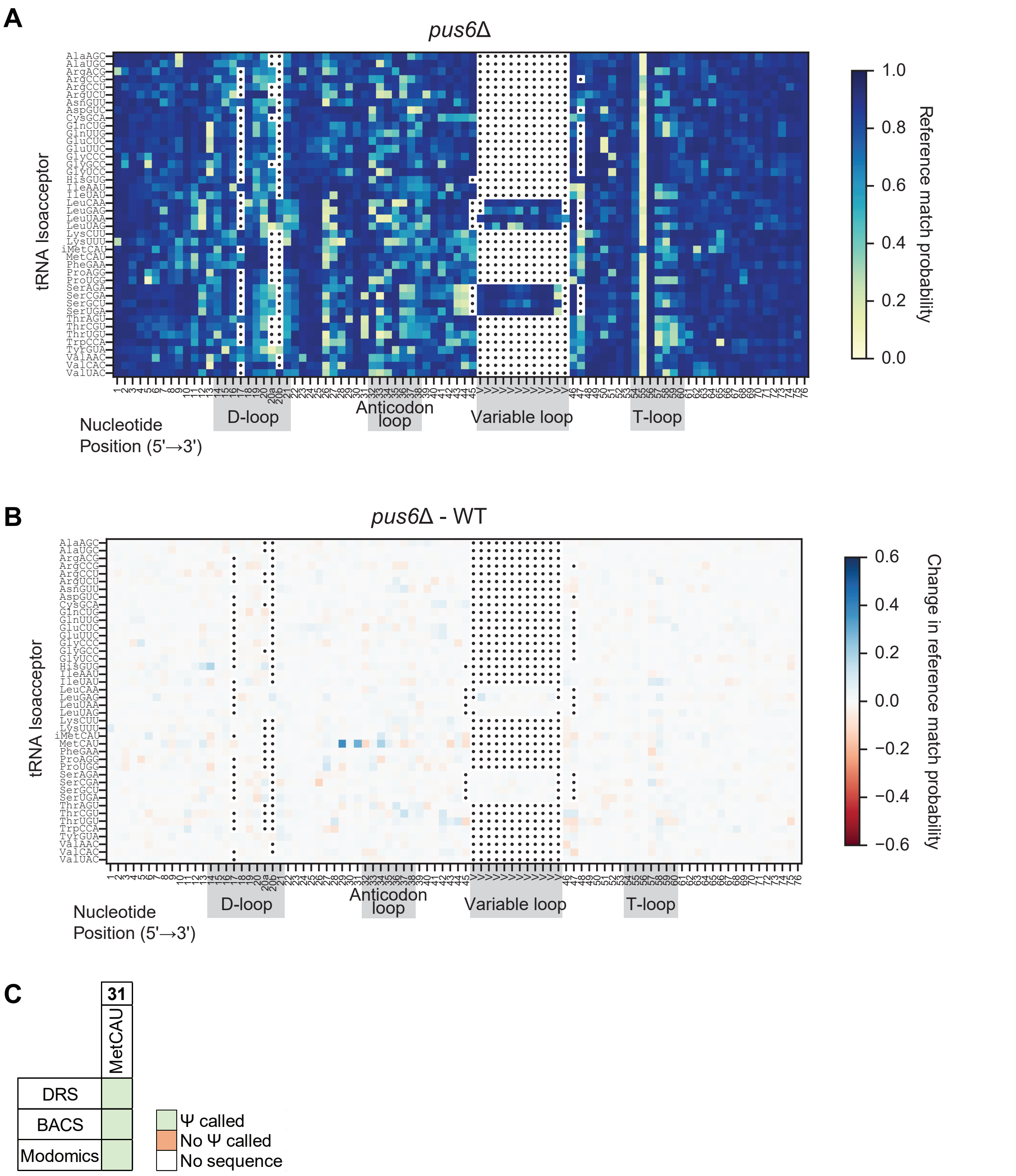


**Figure S8.** Nanopore DRS of budding yeast nuclear-encoded tRNA from a *pus6Δ* strain. (A) Heatmap of Nanopore DRS reference match probabilities of *pus6*Δ yeast nuclear-encoded tRNA. Reference match probability measures the probability that the DRS-called base matches the canonical base in the reference sequence. A low reference match probability (yellow) is suggestive of a chemical base modification. (B) Subtractive heatmap of *pus6*Δ minus wild-type reference match probabilities. A positive change in reference match probability (blue) indicates a decrease in miscalls in the PUS deletion relative to wild-type, while a negative change (red) indicates an increase. (C) Summary of DRS and BACS-detected, and Modomics-annotated pseudouridine sites catalyzed by Pus6, grouped by tRNA position. “No sequence” refers to tRNAs for which no modifications are annotated in Modomics. Here, BACS-called pseudouridines are identified from wild-type data alone, without a corresponding PUS deletion to confirm catalysis.

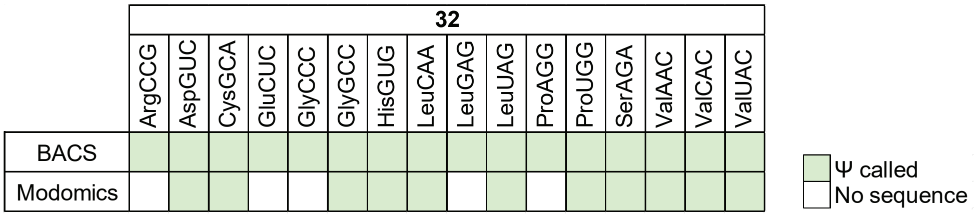


**Figure S9**. BACS analysis of Pus8-catalyzed pseudouridine. Summary of BACS-detected and Modomics-annotated pseudouridine sites catalyzed by Pus8, grouped by tRNA position. “No sequence” refers to tRNAs for which no modifications are annotated in Modomics (Cappannini et al. 2024). Here, BACS-called pseudouridines are identified from wild-type data alone, without a corresponding PUS deletion to confirm catalysis.

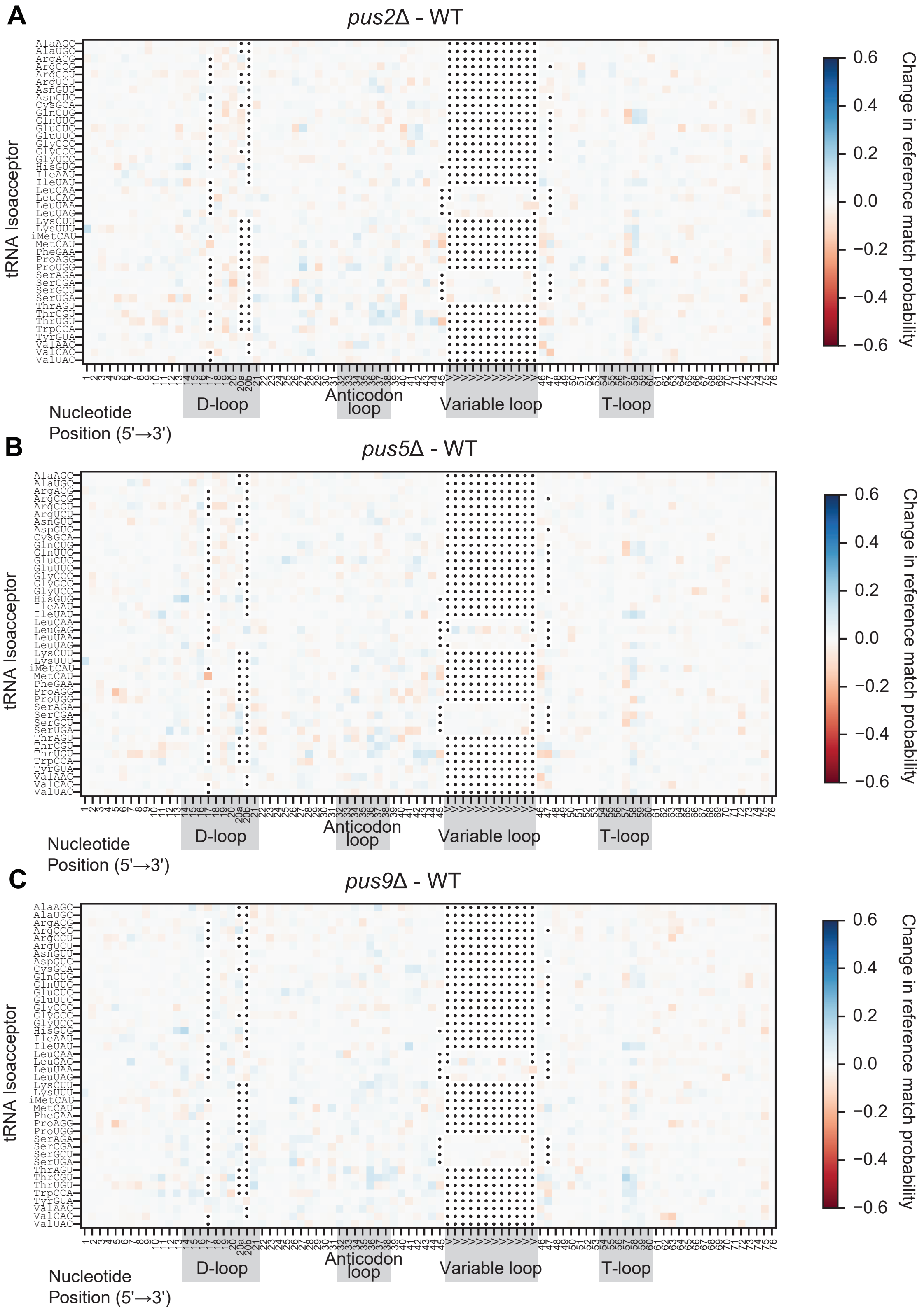
A
pus2Δ - WT
B
pus5Δ - WT
C
pus9Δ - WT
tRNA Isoacceptor
AlaAGC
AlaUGC
ArgACG
ArgCCG
ArgCCU
ArgUCU
AsnGUU
AspGUC
CysGCA
GlnCUG
GlnUUG
GluCUC
GluUUC
GlyCCC
GlyGCC
GlyUCC
HisGUG
IleAAU
IleUAU
LeuCAA
LeuGAG
LeuUAA
LeuUAG
LysCUU
LysUUU
iMetCAU
MetCAU
PheGAA
ProAGG
ProUGG
SerAGA
SerCGA
SerGCU
SerUGA
ThrAGU
ThrCGU
ThrUGU
TrpCCA
TyrGUA
ValAAC
ValCAC
ValUAC
Nucleotide
Position (5'→3')
D-loop
Anticodon loop
Variable loop
T-loop
0.6
0.4
0.2
0.0
−0.2
−0.4
−0.6
Change in reference match probability

**Figure S10**. Nanopore DRS of budding yeast nuclear-encoded tRNA from *pus2*Δ, *pus5*Δ, and *pus9*Δ strains. (A-C) Subtractive heatmaps of reference match probabilities for *pus2*Δ (A), *pus5*Δ (B), and *pus9*Δ (C) relative to wild-type. A positive change in reference match probability (blue) indicates a decrease in miscalls in the PUS deletion relative to wild-type, while a negative change (red) indicates an increase.

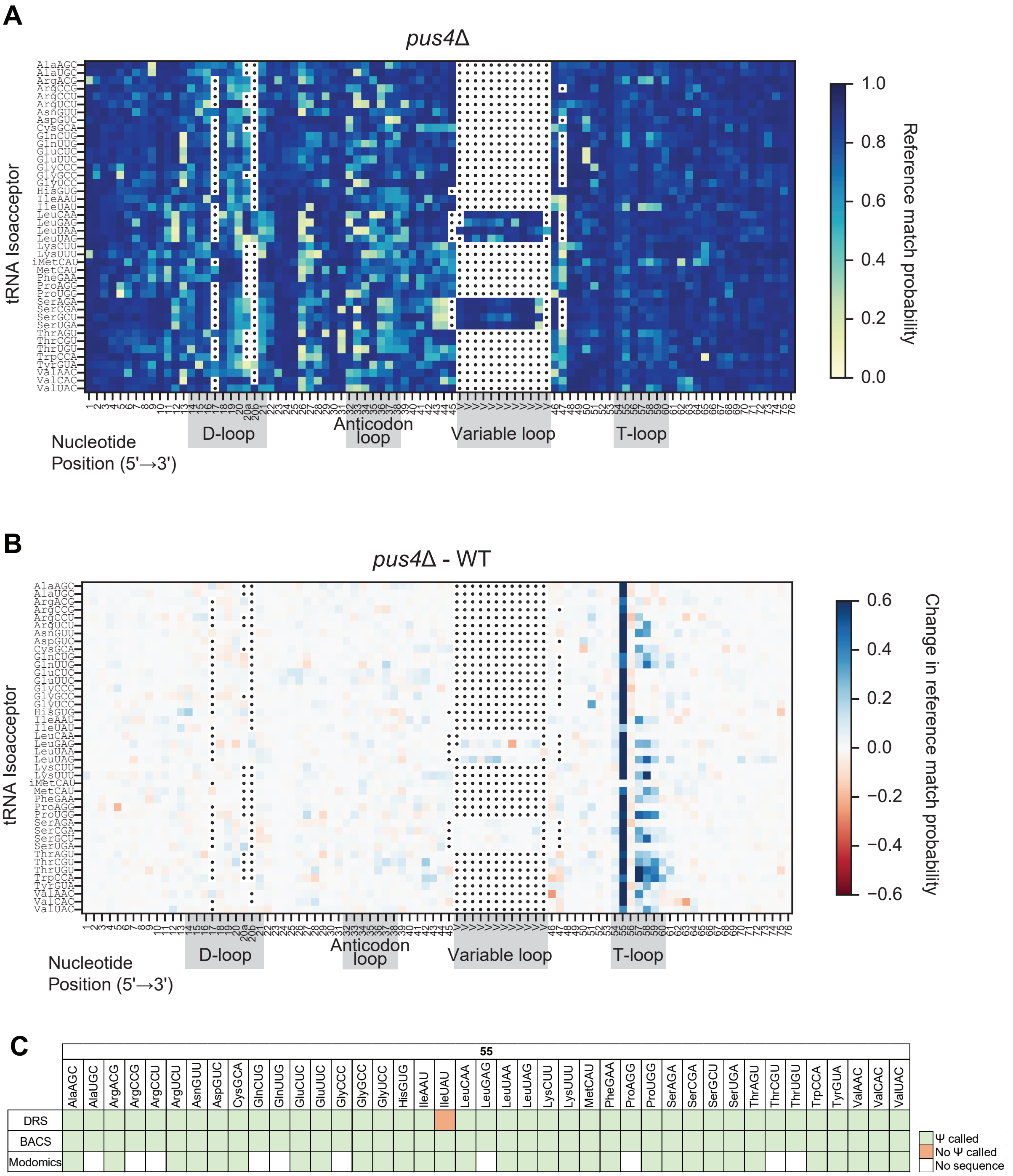


**Figure S11**. Heatmaps of Nanopore DRS data for *pus4*Δ replotted from (Shaw et al. 2024). (A) Heatmap of Nanopore DRS reference match probabilities for *pus4*Δ nuclear-encoded tRNAs, as previously shown in (Shaw et al. 2024). A low reference match probability (yellow) is suggestive of a chemical base modification. (B) Subtractive heatmap comparing reference match probabilities for *pus4*Δ tRNA and wild-type. A positive change in reference match probability (blue) indicates a decrease in miscalls in the PUS deletion relative to wild-type, while a negative change (red) indicates an increase. The decrease in miscalls at position 55 corresponds to loss of pseudouridine in the mutant. The decrease in miscalls at positions

57-59 corresponds to loss of $m^1A_{58}$ with the loss of Pus4. See reference (Shaw et al. 2024) for further discussion. (C) Summary of DRS and BACS-detected, and Modomics-annotated pseudouridine sites catalyzed by Pus4, grouped by tRNA position. For DRS, a green box indicates a change that was dependent on Pus4. For BACS, a green box indicates a pseudouridine called at position 55 in the wild-type data alone, without a corresponding PUS deletion to confirm catalysis. “No sequence” refers to tRNAs for which no modifications are annotated in Modomics. While $tRNA^{Ile(UAU)}$ does contain a pseudouridine sites at position 55 in wild-type based on the BACS data, reanalysis of the DRS data revealed that this signal does not change significantly upon deletion of Pus4. We propose the possibility that the $tRNA^{Ile(UAU)}$ $U_{55}$ is pseudouridylated by a different enzyme in the absence of Pus4, thus retaining a miscall signal in the *pus4*Δ strain.

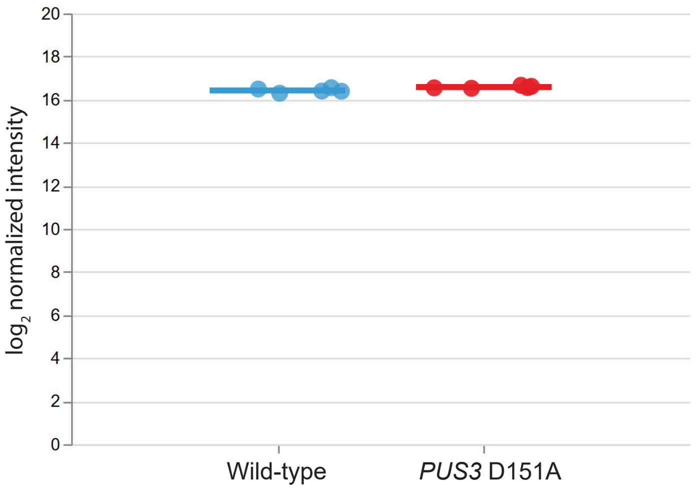


**Figure S12**. DIA mass spectrometry analysis of Pus3 protein abundance in wild-type and *PUS3* D151A cells. Fold change of *PUS3* D151A relative to wild-type is 1.114 ($log_2FC$ = 0.1565; $P$ = 0.034, two-tailed t-test). Each data point is an individual biological replicate. Five biological replicates of wild-type and *PUS3* D151A were grown to exponential phase and harvested. Whole cell lysates were sent to IDeA National Resource for Quantitative Proteomics for data collection and processing.
